# Magnetoencephalography dimensionality reduction informed by dynamic brain states

**DOI:** 10.1101/2024.08.08.607151

**Authors:** Annie E Cathignol, Lionel Kusch, Marianna Angiolelli, Emahnuel Troisi Lopez, Arianna Polverino, Antonella Romano, Giuseppe Sorrentino, Viktor Jirsa, Giovanni Rabuffo, Pierpaolo Sorrentino

**Affiliations:** Faculty of Biology and Medicine, University of Lausanne, 1005 Lausanne, Switzerland; School of Engineering and Management Vaud, HES-SO University of Applied Sciences and Arts Western Switzerland, 1401 Yverdon-les-Bains, Switzerland; Institut de Neurosciences des Systèmes, Aix-Marseille Université, 13005 Marseille, France; Department of Engineering, Università Campus Bio-Medico di Roma, Rome, Italy; Institute of Applied Sciences and Intelligent Systems, National Research Council, Pozzuoli, Italy; ICS Maugeri Hermitage Napoli, Via Miano 69 - 80145; Department of Medical Motor and Wellness Sciences, University of Naples “Parthenope”, Naples, Italy; DiSEGIM, Department of Economics, Law, Cybersecurity, and Sports Sciences, University of Naples Parthenope, Nola, Italy; University of Sassari, Department of Biomedical Sciences, Viale San Pietro, 07100, Sassari, Italy

**Keywords:** Magnetoencephalography, PHATE algorithm, neuronal avalanches, dimensionality reduction, brain dynamics, resting state

## Abstract

Complex spontaneous brain dynamics mirror the large number of interactions taking place among regions, supporting higher functions. Such complexity is manifested in the inter-regional dependencies among signals derived from different brain areas, as observed utilising neuroimaging techniques, like magnetoencephalography. The dynamics of this data produce numerous subsets of active regions at any moment as they evolve. Notably, converging evidence shows that these states can be understood in terms of transient coordinated events that spread across the brain over multiple spatial and temporal scales. Those can be used as a proxy of the “effectiveness” of the dynamics, as they become stereotyped or disorganised in neurological diseases. However, given the high dimensional nature of the data, representing them has been challenging thus far. Dimensionality reduction techniques are typically deployed to describe complex interdependencies and improve their interpretability. However, many dimensionality reduction techniques lose information about the sequence of configurations that took place. Here, we leverage a newly described algorithm, PHATE (Potential of Heat-diffusion for Affinity-based Transition Embedding), specifically designed to preserve the dynamics of the system in the low-dimensional embedding space. We analysed source-reconstructed resting-state magnetoencephalography from 18 healthy subjects to represent the dynamics of the configuration in low-dimensional space. After reduction with PHATE, unsupervised clustering via K-means is applied to identify distinct clusters. The topography of the states is described, and the dynamics are represented as a transition matrix. All the results have been checked against null models, providing a parsimonious account of the large-scale, fast, aperiodic dynamics during resting-state.

## Introduction

Coordination among brain regions is needed to generate appropriate behavioural responses for cognition and to be ready during resting state^1^. The interactions occurring among brain regions are mirrored by statistical correlations between signals representing the activities of the regions. In the healthy brain, the dynamics of these dependencies evolve into complex patterns, continuously rearranging themselves, even during the resting state (i.e. when a subject is not engaged in a specific task).

More specifically, recent evidence suggests that interactions among regions rapidly build and elapse as quickly, generating corresponding transient events of activities that spread over specific spatio-temporal trajectories. Over time, different subsets of brain areas participate in these transient coordinated events^2^. In other words, the interactions at a large scale do not occur continuously but, rather, intermittently^3^. The statistics of these events are generally described by fat-tailed distributions (e.g., power-laws)^4,5^. In the context of critical dynamics, these transient collective fluctuations have been understood as “neural avalanches”^6^. Regardless of whether these transient events genuinely capture the presence of a dynamics operating near a critical point, converging evidence shows that the dynamics of these large-scale states is relevant for cognitive activities, and is altered in case of neurological ailments. From there we can ask ourselves, do patterns of activities spread preferentially along specific spatial trajectories? Furthermore, is there a specific temporal order among such trajectories? Along these lines, several studies focused on the identification of altered brain dynamics via the definitions of microstates^7^. Unlike these studies, we focused here on the dynamics of the aperiodic transient events of activities. We hypothesised that the subset of brain regions recruited by each event are not random but, rather, transient topographies are explored in a structured order with non-trivial transition probabilities. We leveraged source-reconstructed magnetoencephalography data, which offer high spatio-temporal resolution, from 18 healthy young adults, and described the sequence of events by their activation “patterns”, corresponding to the set of brain regions participating in each event. Previous research has highlighted that patterns of neuronal avalanches effectively pinpoint the locations and moments where activities resonate^3^, which is related to the recruitment of brain areas during tasks^8^. Hence, feeding avalanche patterns to a state-of-the-art dimensionality reduction technique might help classify brain states and capture the main features of the high-dimensional avalanche patterns sequence^9,10^. While a plethora of dimensionality reduction techniques exists, they often seem to fail preserving both the local and global similarity of the large-scale brain activities (e.g., with t-SNE^11^). PHATE, on the other hand, preserves these aspects of the data structure in the low-dimensional representation, providing a smoother account of the evolution of the system and, hence, of the sequence of the patterns^12^. Therefore, we utilised this algorithm to reduce the dimensionality of the data. The low-dimensional representations were then grouped into a number of “states” using the k-means algorithm, and the topography of each state was described. Finally, we demonstrated that neither the states we defined nor the transitions between them are expected by chance alone. When the network dynamics are destroyed, e.g., by temporal randomization, the observed patterns also disappear.

## Material and Methods

### Data

#### Participants

A total of eighteen right-handed native Italian speakers participated in the study. Participants were required to meet specific criteria: they should be free from significant medical conditions, substance abuse, and medications that could affect MEG/EEG signals. Additionally, they needed to be free from major systemic, psychiatric, or neurological illnesses and show no evidence of brain damage on routine MRI scans. The study was approved by the Ethics Committee ASL-NA1 centro (Prot.n.93C.E./Reg. n.14-17OSS), and all participants provided written informed consent.

#### MRI Acquisition

Brain images were acquired using a 1.5 Tesla MRI scanner (Signa, GE Healthcare) with a 3D T1-weighted Magnetization-Prepared Gradient-Echo BRAVO sequence. Imaging parameters included a TR of 8.2 ms, TE of 3.1 ms, and TI of 450 ms, with a voxel size of 1 × 1 × 1 mm³ and a 50% partition overlap across 324 sagittal slices covering the entire brain.

#### MEG Acquisition

Magnetoencephalographic data were collected using a 163-magnetometer MEG system in a magnetically shielded room (AtB Biomag UG, Ulm, Germany). Preprocessing followed established protocols. Four coils and four reference points (nasion, right and left pre-auricular points, and apex) were digitised using the Fastrak (Polhemus®) prior to acquisition. Brain activity was recorded for ∼7 minutes with eyes closed, divided in two halves to reduce drowsiness. The head position was measured at the start of each segment. Data were sampled at 1024 Hz and filtered with a 4th order Butterworth band-pass filter between 0.5 and 48 Hz. ECG and EOG were recorded during acquisition. Matlab 2019a and the Fieldtrip toolbox 2014 were used for preprocessing.

### Data processing

#### Preprocessing

To reduce environmental noise, principal component analysis (PCA) was employed. Noisy channels were manually removed through visual inspection by an experienced rater. Supervised independent component analysis (ICA) was utilised to remove ECG and EOG artefacts from the MEG signals. Trials free from artefacts or excessive noise were selected for further analysis.

#### Source Reconstruction

The MEG data were co-registered with native MRI scans^13^. A modified spherical conductor model served as the forward model. Voxels in the MRI were labelled according to the Automated Anatomical Labelling (AAL) atlas, focusing on 78 cortical regions. A linearly constrained minimum variance beamformer computed 78 time series, each corresponding to one region of interest, at a sampling frequency of 1024 Hz. An experienced rater visually inspected the reconstructed sources. The source-reconstructed signals were then downsampled to 256 Hz.

#### Avalanche pattern

For each subject, each source-reconstructed MEG timeserie was standardised (z-scored) and binarized based on a threshold fixed at 3 SD, following the same value chosen in a previously cited study^8^.

Deviations above the threshold, marked by ones in the binarized signals, tend to co-occur across subsets of brain regions, which suggests the presence of spatiotemporal correlations and collective behaviours operating across different cortical areas^14^.

Based on the binarised signals, we defined neuronal avalanches as starting when at least one region is above threshold, and ending when no region is above threshold^8^. Then, for each avalanche, we established an *avalanche pattern* as a vector of size *N*, with *N* representing the number of brain regions, containing 1 if a region was recruited during the avalanche (at any point, and for any duration), and 0 otherwise. In other words, we analysed the spatial structure of each avalanche, while disregarding the internal structure.

We then concatenated all the unique avalanche patterns, and used these as the input for the dimensionality reduction algorithm, PHATE. Figure <u>1</u> and <u>2</u> provide respectively an overview of the pipeline and an overview of the various stages of the input data in a low-dimensional reduction with PCA.

**Fig 1.**
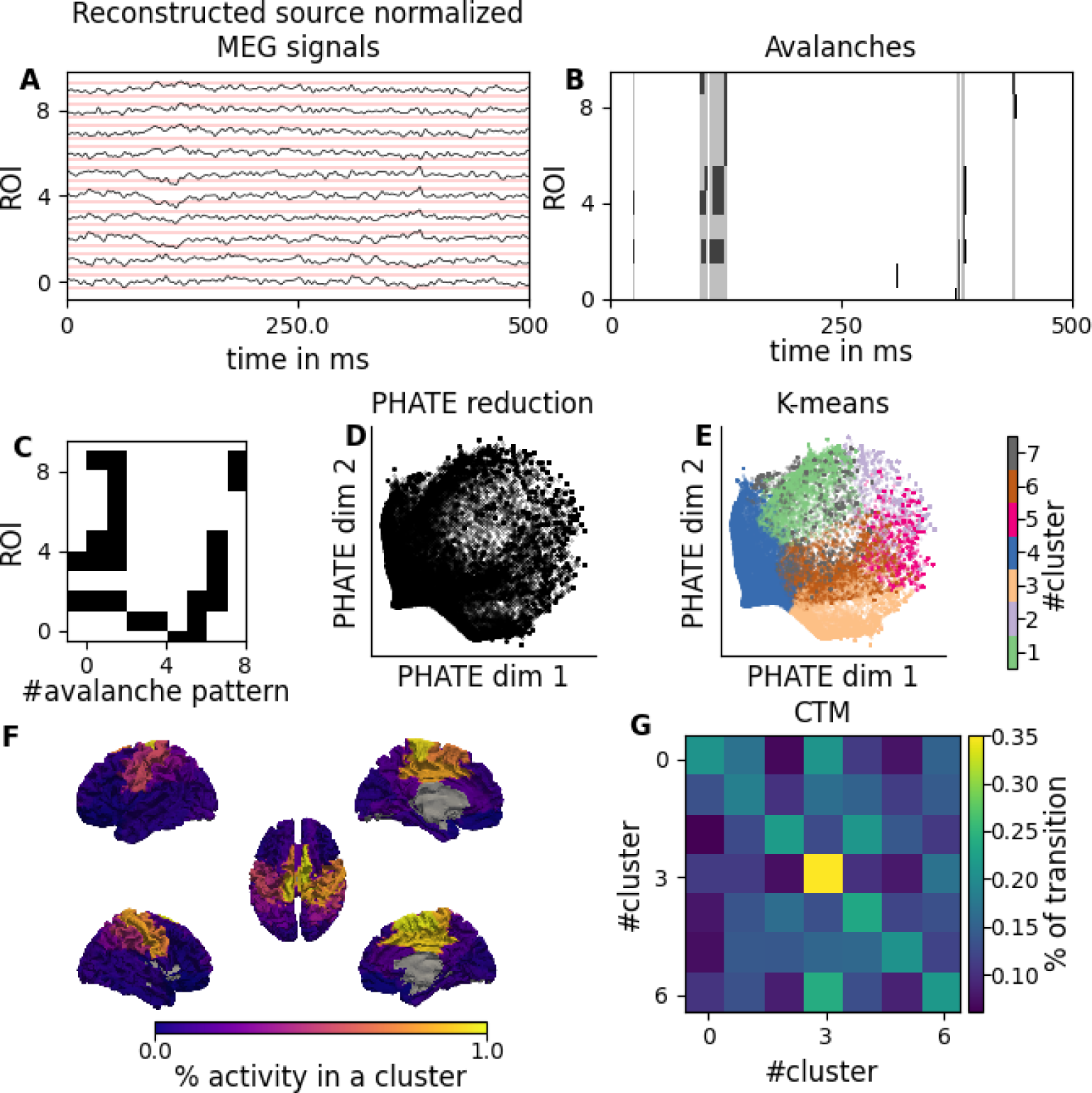
Overall Pipeline steps. Magnetoencephalography (MEG) source-reconstructed signals are used to detect neural activity (**A**). After normalising each signal and z-scoring them, transient coordinated events of brain activity across multiple channels (i.e., neuronal avalanches) are identified (**B**). Each avalanche is then vectorized to define an "avalanche pattern," capturing the active regions during the event and encapsulating the spatial and temporal information about brain activity (**C**). The dimensionality of the data is reduced using a technique called PHATE (**D**) (Potential of Heat-diffusion for Affinity-based Trajectory Embedding), which helps visualise and interpret high-dimensional data in a low-dimensional space. Finally, K-means clustering is applied to the primary PHATE components to extract the final clusters (**E**), resulting in the identification of seven clusters characterised by their average activity vector (for example, the cluster 6) (**F**). The probability matrix of changing clusters (**G**).

**Fig 2.**
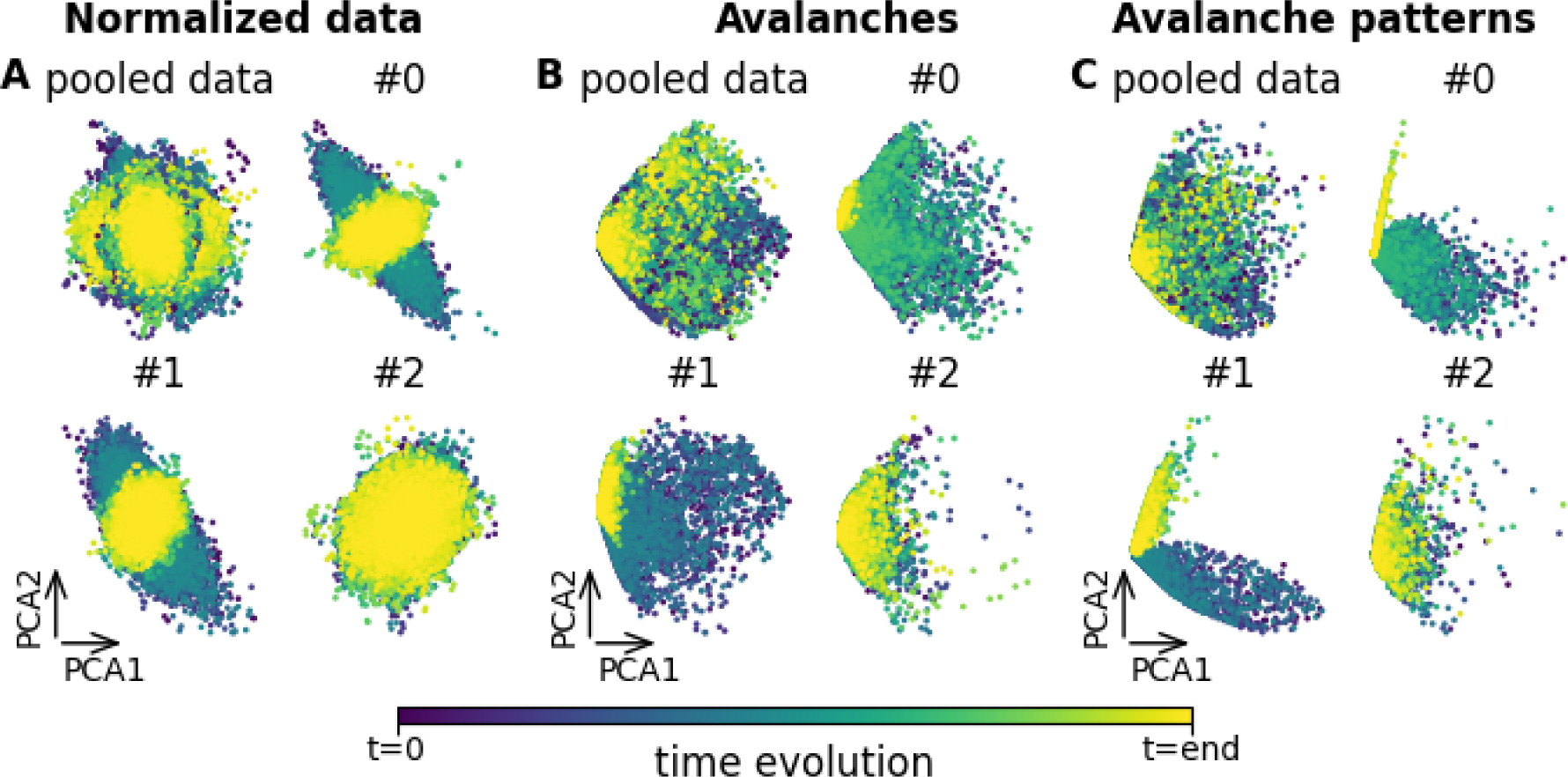
Low-dimensional reduction using PCA. The picture shows the time evolution of the system by subject in PCA space for the different steps of the pipeline and the low-dimensional reduction using PCA detection starting from normalised data (**A**), avalanche activity (**B**) and avalanche patterns (**C)**. In each subplot, each dot represents the evolution of the whole system brain state at each instant along the first (PCA1) and second (PCA2) dimension. (the colour gradient from dark purple to yellow indicates the time evolution from the beginning to the end of the recording) for pooled data and the subject 0, 1 and 2 as examples. We notice how focusing on avalanches most details are discarded from the data, but this reveals an underlying structure that it is already possible to capture using PCA.

#### Potential of Heat-diffusion for Affinity-based Transition Embedding algorithm: PHATE

PHATE (Potential of Heat-diffusion for Affinity-based Transition Embedding) is an algorithm designed to visualise multidimensional data by a nonlinear dimensionality reduction that preserves, in the low dimensions, the local and global structure of the data. This algorithm is described in detail in Moon et al. 2019^12^. In a nutshell, the first step is the application of a Principal Component Analysis (PCA) where only the most informative components are retained. In this study, the first five components were identified as the most significant based on the explained variance and their prominence in the cumulative variance plot (see <u>Sensibility analysis</u> for more details, as well as supplementary figure <u>SP1</u> and <u>SP2</u>). The procedure to preserve local and global structures in low dimensions relies on calculating the distances between each avalanche pattern (a point in high dimensional space) using cosine similarity (figure <u>SP3</u>). These distances are then transformed into a Markov-normalised affinity matrix using a kernel function. For our specific application, the kernel was configured to consider the five nearest neighbours, with an ɑ-decay of 1 (see <u>below: Assessment of the ɑ-decay parameter for the PHATE algorithm</u>). Based on this, a diffusion probability is calculated and used directly to embed the cosine distance with multidimensional scaling into a three-dimensional space.

#### Unsupervised Machine Learning: K-means clustering

After the dimensionality reduction via PHATE (or via a simple PCA as for figure <u>3</u>), we classified the points in low dimensions using unsupervised K-means clustering^15^. This algorithm operates iteratively to group data into clusters. The first iteration randomly initiates a number (here *k*=7) of cluster centroids. Each point in the training dataset is then assigned to the nearest centroid based on euclidean distance (the method for selecting the number of centroids, *k*, is detailed in the <u>Sensibility analysis</u> section that follows). In the subsequent steps, each centroid is moved to the mean of the points assigned to it. After one iteration is finished, the next iteration will again assign each training sample to the closest cluster centroid and move those centroids again according to the mean of the points. Either a maximum number of iterations is fixed, or the algorithm will continue until the change of position of the centroids is under a predetermined “tolerated” value^16^.

**Fig 3.**
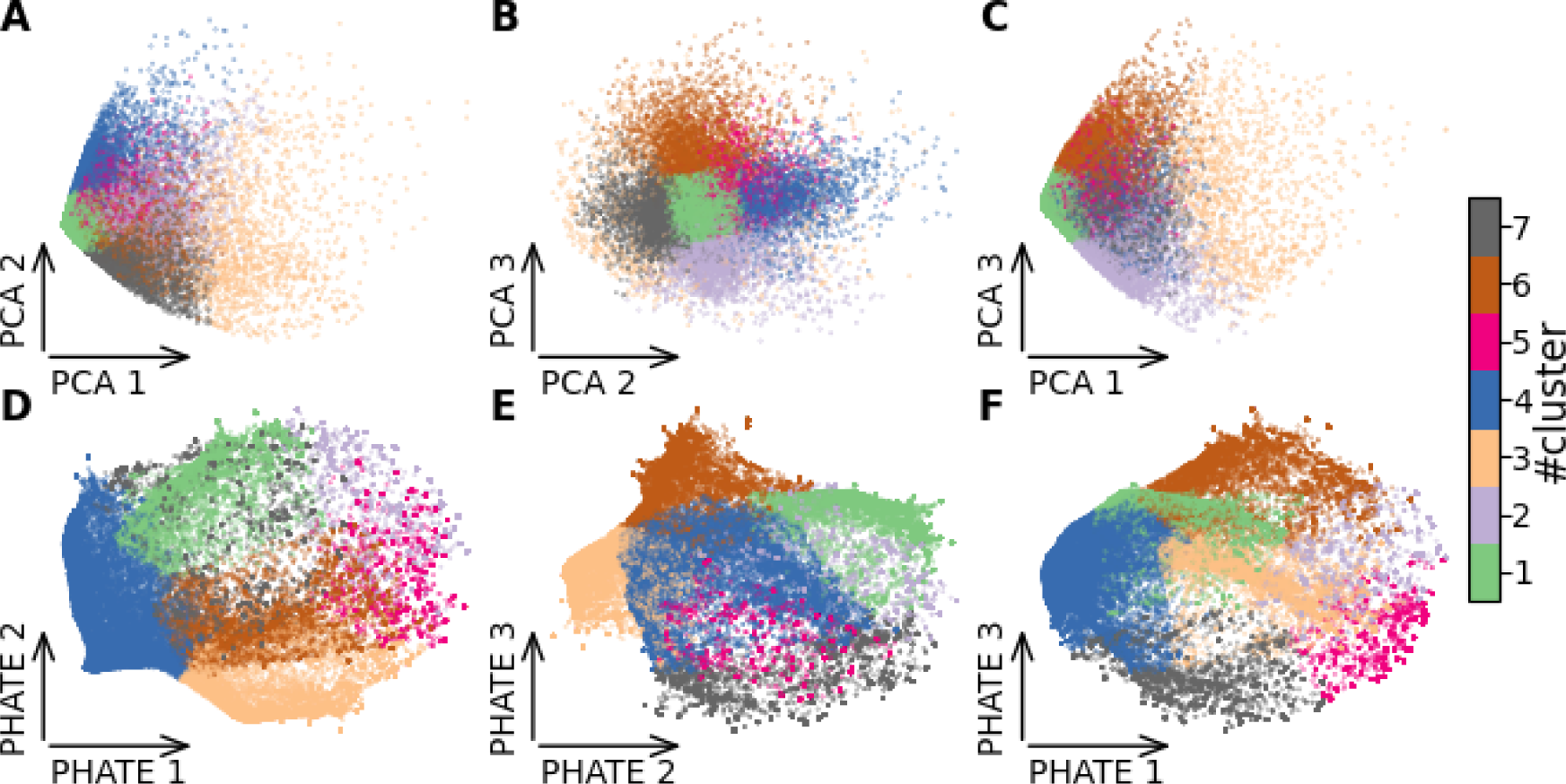
Comparison of low dimensions of PCA and PHATE. The process of dimensionality reduction is done using the PHATE (Potential of Heat-diffusion for Affinity-based Trajectory Embedding) algorithm. Initially, PCA is used to reduce the data into five principal components, the three firsts are shown in the scatter plots (PC1, PC2, PC3 in panels **A**, **B** and **C**). The data points are then further processed through PHATE, which reveals a more nuanced structure, effectively separating the clusters based on intrinsic data properties. This transformation is visualised in a two-dimensional scatter plot of the first three axes (PHATE 1, PHATE 2 and PHATE 3 in panels **D**, **E** and **F**). It highlights distinct clusters, which are subsequently categorised using K-means clustering for 7 clusters (indicated by various colours). For all panels, each point represents an avalanche pattern sorted on the axes of the primary components (PCA or PHATE).

#### Transition between clusters

Based on the clusters identified by the k-means algorithm, we labelled each avalanche pattern and computed the transition probability matrix for each subject, capturing the transitions between clusters (figure <u>4.H</u>). Additionally, we computed a global transition matrix that aggregates the avalanche patterns across all subjects. Notice that the latter includes some “false transitions” due to the concatenation of avalanche patterns belonging to different subjects. However, due to the low number of “false transitions”, this should not bias the estimated probability.

**Fig 4.**
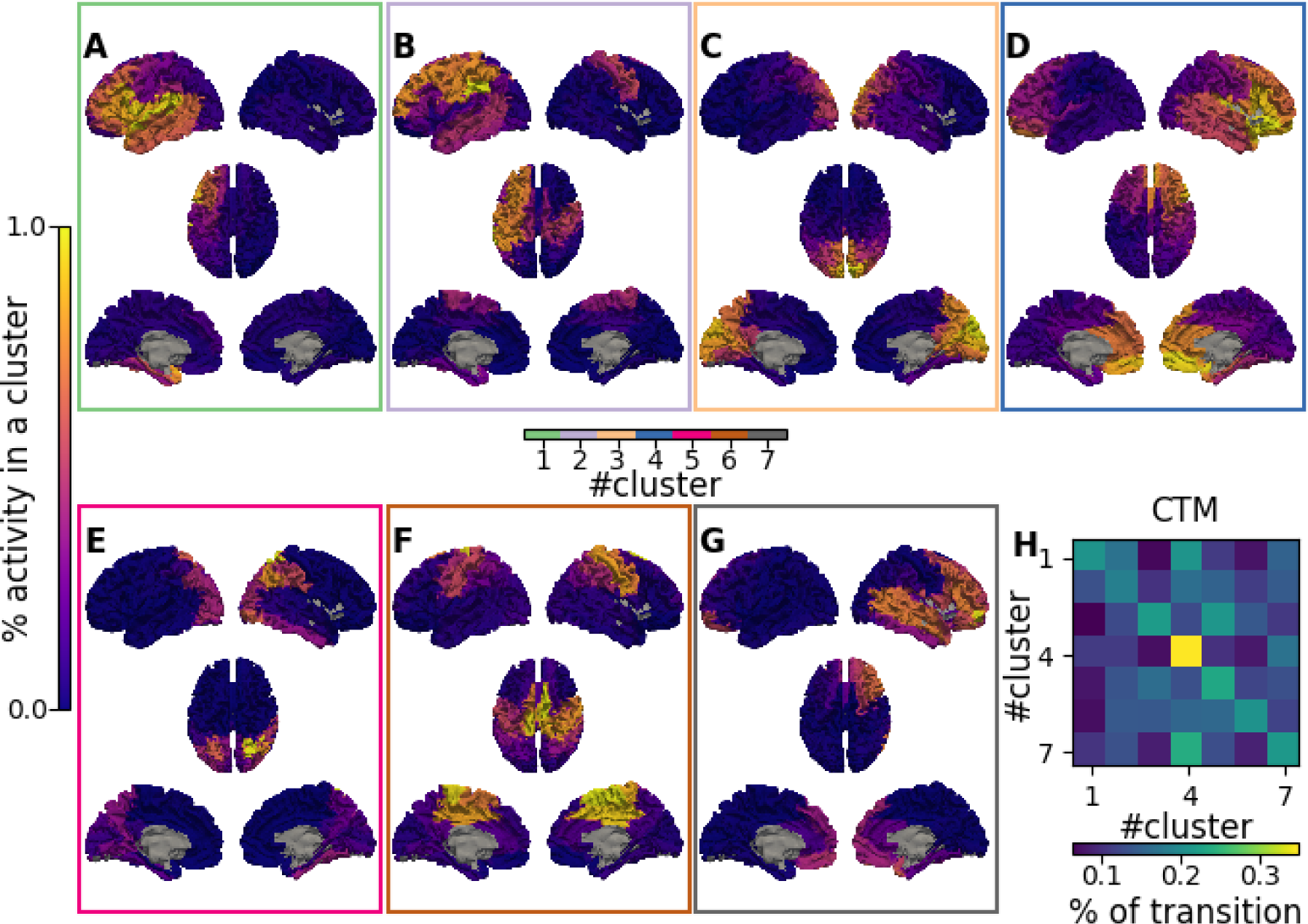
Output of the Pipeline on brain images and matrices. The pipeline is applied on the avalanches patterns to reduce the dimension with PHATE and then cluster each avalanches with K-means algorithms. The result of the pipeline is the average activity of each cluster of our output projected on brain images (**A**, **B**, **C**, **D**, **E**, **F**, **G**). More precisely, it shows the projection on brain images of the statistical activation (yellow) or inactivation (dark blue) of regions for each cluster. The second output of the cluster is (**J**) the probability of transition between each cluster, i.e., the transition matrix.

### Sensibility analysis

Sensibility analysis was applied to evaluate the uncertainty about the most important parameters used in our pipeline: the number of components in the PCA and the ɑ-decay for the PHATE algorithm, as well as the number of clusters for the k-means.

#### Assessment of the number of components in the PCA of PHATE algorithm

To determine the number *C* of components in the first step of PHATE, we plotted the PCA explained variance relative to the number of components. For this, the evolution of the cumulative explained variance was plotted against the number of components of the PCA (see supplementary figure <u>SP1</u> and <u>SP2</u>). The optimal number of components was selected such that no visual significant difference in explained variance was observed by using *C* or *C+1* components, ensuring that adding more than 5 components does not significantly improve the explained variance.

#### Assessment of the ɑ-decay parameter for the PHATE algorithm

In order to fix the PHATE parameter ɑ-decay of 1, we first used empirical tests, trying to reduce it in order to capture the smallest local structures. However, a minimal value of the decay can be estimated by calculating the maximal distances between neighbours provided by the K-nearest neighbour algorithm (figure <u>SP3</u>). This estimation is based on the calculation of the kernel which requires to provide a significatif weight with the desired number of neighbours.

#### Assessment of the number of clusters in the K-mean clustering (for different kernels of PHATE)

We assessed the number of clusters by looking at three different graphical methods: the elbow measure, the silhouette method and the gap statistic ^17^.

The elbow method is based on a graph showing the within-cluster-sum-of-square (WCSS) values for different numbers of centroids (*k*). We described the gain in explained variance as a function of the number of clusters (figure <u>SP1</u>). The point of flexion provides an estimate for the optimal number of clusters.

The silhouette method (figure <u>SP1</u>) is based on plotting the average silhouettes of our observations for various numbers of centroids. A good clustering is indicated by a high average silhouette. The optimal k is the one that maximises the average silhouette.

The gap statistic method (figure <u>SP1</u>) is based on comparing the difference, i.e., *gap*, between the expected total intra-cluster variation and the observed one for a specific *k*. This is then repeated for different values of *k* and plotted as a function of the number of clusters. The higher the gap, the more a particular number of clusters provides a parsimonious account of the variance in the data.

To visualise the topography of the clusters, we computed how much each region is recruited in the patterns belonging to each cluster (summing the number of times each region is active in patterns belonging to a given cluster). This way, we describe cluster-specific topographies.

### Statistical validation

The robustness of the results was evaluated by comparing our observations to null models. In particular, we analysed the robustness of 1) the activation pattern of each cluster (figure <u>5.A</u>, <u>5.B</u>, <u>5.C</u>), 2) the transition prediction between clusters (figure <u>5.D</u>, <u>5.E</u>, <u>5.F</u>, <u>5.G</u>, <u>5.H</u>), and 3) the dimensionality reduction (figure <u>5.J</u>, <u>5.K</u>, <u>5.L</u>).

**Fig 5.**
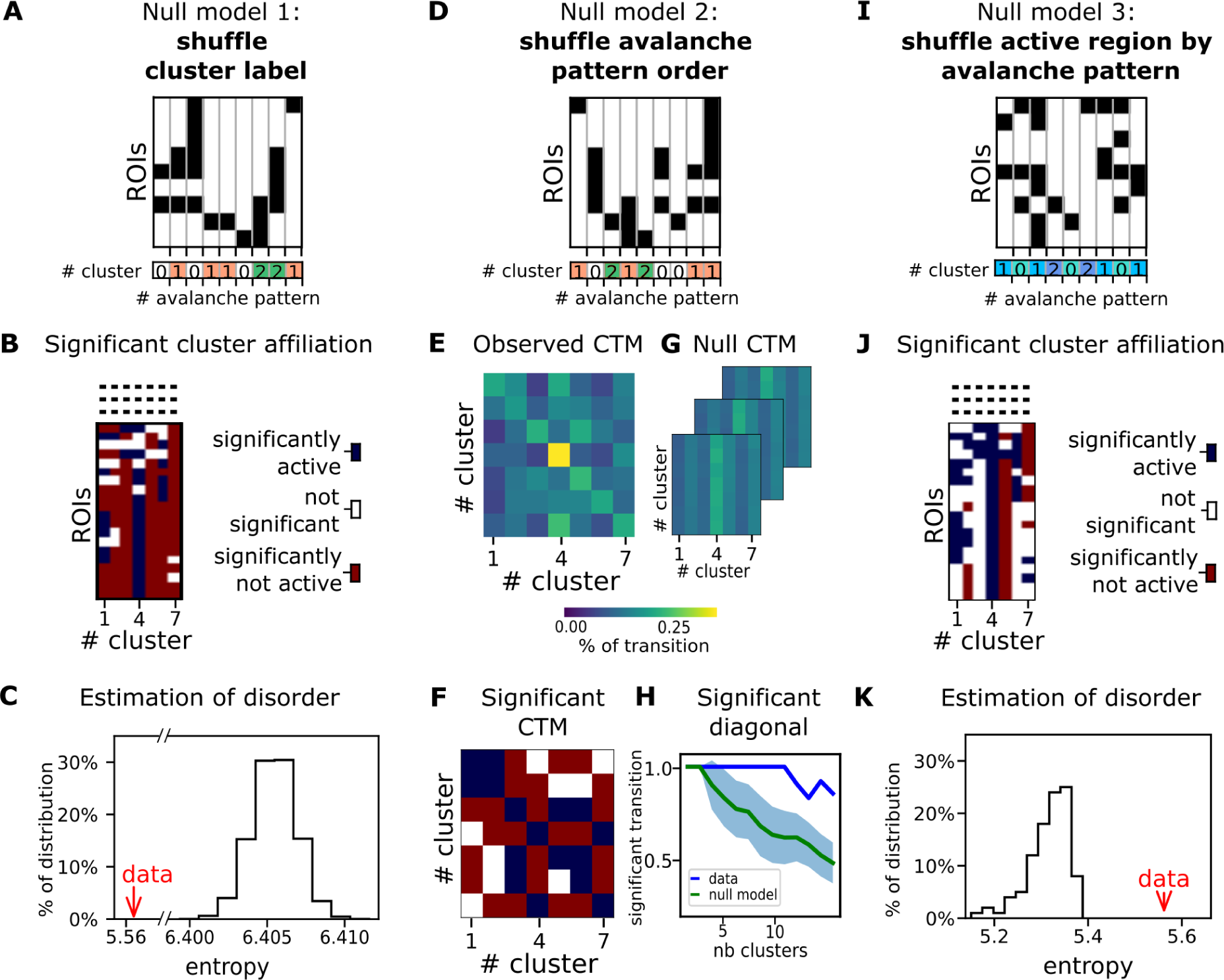
Robustness analyses. The figure highlights the 3 methods to check the robustness of our findings, based on null models. The first solution (**null model 1**) mixes cluster labels (**A**) and observes the entropy distribution (**C**) resulting from such shuffling (in black) versus our original one (red arrow). The panels **B** (as the following panel **J)** displays the partial vectors of all clusters (the full vectors are displayed in figure <u>SP4</u>). In other words, they represent the sum of avalanches belonging to the cluster, resulting from the association of each avalanche to a specific cluster. More specifically, those panels show the significance of the probabilities of an avalanche activation (red), not activation (blue) or at basal state (white) to be part of a cluster. The second null model (**null model 2**) is based on shuffling the avalanche patterns’ order (**D**) to test the robustness of the transition. Matrices **E**, **F** and **G** display respectively the probabilities of transitions between each cluster for our original data, the significance of the probabilities of a cluster and the shuffled ones. The probability of transition highlights a higher transition probability within clusters, but still lower than chance level (**H**). The last one (**null model 3**) is to shuffle each active region in each avalanche pattern (**I**) and check the robustness of the dimensional reduction computing the entropy distribution of the cluster label shuffled output (black distribution) and entropy measure of our original output (red arrow) (**K**). This tests the robustness of the average vectors of each cluster, in new dimensions (indicated by the colour blue).

The null model hypothesis suggests that any observed differences in the studied characteristic from a dataset are attributable to chance. Hence, for each robustness analysis (point 1), 2) or 3)), we mixed the data such that the studied feature was found random; each analysis requires a different way of randomising the data, hence, results in a different “null dataset”. We then compared the characteristic between the existing dataset and the random ones, demonstrating if a statistical property in the data was to be expected by chance.

#### Robustness of the activation pattern of each cluster

First, based on the null model hypothesis, we wanted to test if the clusters obtained by our algorithm were informative and not random. As a result of the algorithm (PHATE and k-means), each avalanche pattern was associated with a cluster. From this association, as explained above, we described the activity pattern of each cluster, i.e., a vector containing the average activation of each brain region (e.g., brain plots in figure <u>4</u>). To check that the clusters were non-random, we computed the entropy of the cluster matrix, i.e. the matrix obtained by concatenating all the patterns (regardless of the clustering). We compare the empirical entropy to the ones observed in the null cluster patterns. The entropy of point x, H(x), was calculated using Shannon’s equation^18^:

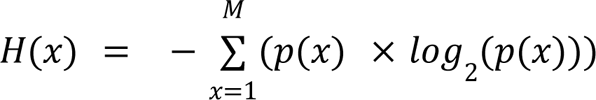

where *p(x)* is the percentage activation of each brain region in a cluster, and *M* is the length of the concatenated cluster vectors.

This way, we obtained the distribution of the entropies of the null data, against which we compared the entropy of the clusters we observed from non-random labelling (figure <u>5.C</u>).

Comparing our original output with the null output we also verified, for each brain region in each cluster, if any given region was more (less) recruited in the patterns belonging to a particular cluster (figure <u>5.B</u> and <u>SP4</u>). We did this by comparing the probability of any region being recruited in the observed clusters to the probability of the same regions being recruited in a pattern obtained after the shuffling. We defined the significance threshold as 0.05.

#### Robustness of transition prediction between clusters

For each subject, we tested whether the transitions between the clusters were statistically significant. This involved generating shuffled sequences of each subject’s data to serve as a baseline for comparison. Transition predictions between clusters for each subject were then evaluated against these shuffled sequences (figure <u>SP4</u>). The transition probabilities of the original sequence were compared to those derived from the corresponding shuffled sequences, computing the correlation between them, and yielding a p-value for each transition probability (figure <u>SP6</u>). Looking at the resulting p-value matrix for each subject, we could identify statistically significant transitions (both occurring more or less often than what is expected by chance alone) at a significance level of 0.05. The transition sequences per subject also enabled us to evaluate inter-subjects variability (e.g., mean, standard deviation and coefficient of variation between the transitions).

Finally, we used the same method described in the paragraph above to determine whether the transitions between clusters were non-random across subjects, as predicted by the null model. This analysis involved comparing the transition probabilities from the concatenated sequences against those from shuffled sequences, mirroring the approach used for individual subject data (figure <u>5.F</u>). We calculated the p-values for each transition probability in the concatenated sequences to assess statistical significance (figure <u>5.H</u>). Similar to the individual analyses, significant transitions were identified based on a threshold of 0.05.

This way, we could estimate to what extent the pattern of the transitions is shared among subjects. Indeed, if the prediction of transition is robust between clusters, an additional analysis can be done to identify consistent cluster transitions across individuals. This consistency would suggest that rather than being the result of individual variances or random variation, the observed transitions are most likely indicative of the population under study.

#### Robustness of dimensionality reduction

To quantify the effectiveness of the dimensionality reduction, we used again the null model hypothesis, this time shuffling active and non-active regions within each pattern. To this end, we shuffled the labels of all regions —both active and inactive— within the input data for each avalanche prior to feeding it into PHATE, for dimensionality reduction, and k-means, for clustering. Subsequently, we assessed the robustness of the dimensionality reduction by comparing the entropy of the low dimensional output obtained from the original data against the entropy distribution observed when starting from the shuffled data (figure <u>5.L</u>). As described above, we verified, for each region and for each cluster, the probability of the activation or deactivation of the region if the null model hypothesis was true, providing an estimate of the confidence that the activations of any regions that are included in a given pattern are non-random (figure <u>5.K</u> and <u>SP4</u>). We considered a statistical significance threshold of 0.2. The reduction of precision is due to the fact that the instances of the different clusters are not homogeneous (that is, some clusters are present more often than others). As a consequence, the statistical power and the detectable effect size change across patterns.

## Results

### Combining neuronal avalanches and dimensionality reduction to characterise brain network dynamics

Starting from the source reconstructed MEG signals (figure <u>1.A</u>), we identified the sequence of neuronal avalanches through a binarization process; (figure 1.B; see <u>Material and Methods</u>), and defined for each avalanche the set of regions that were recruited as an *avalanche pattern* (figure 1.C). Then, we fed the sequence of avalanche patterns to the PHATE algorithm to reduce their dimensionality (figure 1.D). This way, we preserved the spatial information about the patterns as well as their evolution over time. Applying the K-means algorithm to the first components of PHATE (figure <u>1.E</u>) allowed us to cluster the avalanche patterns into self-similar groups. Averaging the avalanche patterns within each cluster resulted in characteristic brain networks with nontrivial topography (e.g., figure <u>1.F</u>). Studying the original sequence of avalanche patterns, and labelling the patterns according to the corresponding cluster, we extracted the transition probability matrix, or Cluster Transition Matrix (CTM). We have also repeated the analysis using PCA as a dimensionality reduction technique, to compare the performance with that of PHATE. Note that neuronal avalanches are based on a binarization algorithm that greatly reduce the amount of information in the data: signals can only have 0 (‘inactive’) and 1 (‘active’) values, and the threshold for binarization is set in such a way that only a small percentage of time points (e.g., 5%) is considered as active. Nonetheless, focusing on neuronal avalanches prior to applying dimensionality reduction techniques (i.e., PHATE and PCA) allowed us to extract valuable information regarding the topography and dynamics of brain states, while reducing the noise in brain data.

### Focusing on neuronal avalanches reveals hidden low-dimensional structure

To analyse the advantages of reducing data informed by neuronal avalanche algorithm, we analysed the performance of dimensionality reduction techniques when applied to the raw data (figure <u>1.A</u>), to the avalanche activity (i.e., without averaging the activity within each avalanche; figure <u>1.B</u>), or to the avalanche patterns (figure <u>1.C</u>). We start by focusing on the PCA dimensionality reduction, projecting the brain activity (raw signals, avalanche activity, or avalanche patterns) onto the first two PCA components (figure <u>2</u>). Feeding the avalanche patterns to the PCA (figure <u>2.C</u>) highlights a more rich low-dimensional data structure as compared to the normalised raw data (figure <u>2.A</u>) or to the avalanche activity (figure <u>2.B</u>). In general, using the avalanche activity and the avalanches patterns (figure <u>2.B</u> and <u>2.C</u>, respectively) improves the separation of the data points as compared to the normalised data (figure <u>2.A</u>). Furthermore, as observed in figure <u>SP7</u>, the use of avalanches highlights a common structure, removing the specificity of subjects and highlighting common structures. In the supplementary figures, we report the data reduction starting from the reconstructed z-scored MEG source signals (figure <u>SP8</u>), from avalanches (figure <u>SP9</u>), and from the avalanches patterns (figure <u>SP10</u>) for all the subjects, and for each of the 18 subjects individually. This suggests that the avalanche analysis can enhance the interpretability, as well as the visualisation of the underlying neuronal dynamics as compared to using normalised raw data. In other words, focusing on the avalanche patterns provides a theoretically-informed, yet data-driven way to select only a handful of points in time and space which are most significant. Similar trends were observed employing PHATE to reduce the dimensionality of the data (figure <u>3</u>; see also figure <u>SP11</u> where different pipelines were used).

### PHATE dimensionality reduction outperforms PCA on brain state identification

As explained in the methods, PHATE aims at preserving both the details and the high-level structures that are present in the high-dimensional representation of the data^12^. The PHATE algorithm entails a preliminary dimensionality reduction step based on PCA, a heat-diffusion processes to help denoising the data, while preserving the manifold distances in the lower-dimensional embedding space.

Here, we applied PHATE or PCA to subject-specific avalanches patterns, and extracted the main avalanche patterns clusters using K-means (as in figures <u>1.D</u>, <u>1.E</u>). Compared to PCA, PHATE retrieved clusters that are better separated, displaying a clearer three-dimensional structure (figure <u>3</u>). In fact, PCA fails to show a clearly identifiable number of clusters. The PHATE algorithm detected seven clusters, identifying more patterns in the data and, as a result, helping the clusterization.

Supplementary figure <u>SP1</u> displays the elbow, silhouette, and gap statistics graphs for both the PHATE and the PCA reductions.

In addition, the cumulative activations of each region, indicates that no specific region seems to drive the classification (supplementary figure <u>SP12</u>). This would suggest that the clustering captures genuinely multivariate patterns.

### Identification of brain state topographies using PHATE dimensionality reduction

We visualised how often each brain region was recruited in each cluster, providing a topographical account of the clusters (figure <u>1.F</u> - see <u>method</u> for more information). Figure <u>4</u>, shows that the cluster-specific topography varies sharply, with each cluster showing an interpretable pattern of regions that are preferentially recruited. On the one hand, in clusters 3 (<u>4.C</u>) and 5 (<u>4.E</u>), the occipital regions are the most active. Cluster 2 (<u>4.B</u>) and 4 (<u>4.D</u>), on the other hand, display symmetrical activities in frontal regions, with minor involvement of the contralateral fronto-parietal regions. The Frontotemporal regions are symmetrically involved in clusters 1 (<u>4.A</u>) and 7 (<u>4.G</u>), while cluster 6 (<u>4.F</u>) hinges around the sensorimotor regions. As seen, despite the clustering being data-driven, the clusters are reminiscent of known anatomical/functional patterns, supporting the validity of the cluster-specific topographies.

#### Large-scale dynamics

##### Identification of brain state topographies

Studying the transition matrix between clusters (CTM; figure <u>1.G</u>) we show that, despite neuronal avalanches being separated by long inter-avalanche intervals, their sequence is far from being a random process. Much like an earthquake has a higher probability of being followed by aftershocks, we observe that the transition to the same cluster (i.e., as if the cluster is recruited multiple consecutive times), is the most probable transition, as shown by the high values of the main diagonal in figure <u>4.H</u> (or figure <u>1.G</u>). This can be seen as evidence that the system possesses a memory about the previously recruited pattern. The clusters could thus be considered quasi-stable in time. However, some inter-cluster transitions display a high probability of occurring. Overall, most transitions are occurring with a probability higher than chance-level, as shown in figure <u>SP13</u>. Similarly, supplementary figure <u>SP14</u> displays the mean, standard deviation and coefficient of variation of the transitions between clusters for all subjects. We observed that the mean of the diagonal remains quite stable, around 0.2, with little variability except for cluster 3, which has higher mean and higher variability, and cluster 1, which shows high variability. Other features could be observed as well: cluster 6 has a tendency to transit to cluster 3, while the latter has a tendency to be recruited multiple times or to transit to cluster 6. Finally, cluster 1 has a tendency to transit to cluster 2 and 3, while cluster 2 has a tendency to transit to cluster 1.

However, when displayed across subjects (figure <u>SP6</u>), the significance of the transition probability between clusters highlights a high variability. This observation, already visible in the matrices of transition probabilities across subjects (figure <u>SP5</u>), could be explained by the short span of the recordings. However, figure <u>SP15</u> shows that the same transitions (i.e. between the same clusters) occur above/below chance across subjects. In other words, the dynamics we observed is consistent across subjects, despite the dimensionality reduction being performed independently for each.

##### Sensibility analysis and statistical validation

The sensitivity analysis, which has been carried out to test the robustness of our results (see <u>Material and Methods</u>) involved the evaluation of the number of components for the PCA within the PHATE algorithm,and the optimal number of clusters for the K-means clustering. To this end, we considered various parameters and statistical or visual measures, such as the explained variance and the gap statistics.

### Assessment of the number of components in the PCA of PHATE algorithm

The assessment of the number of components in the PCA, which is carried out as the first step of the PHATE algorithm (figure <u>SP1</u> and <u>SP2</u>) showed a marked drop until the fifth component (included). In fact, the explained variance went from about 0.125 with one component, to roughly 0.35 with five components. When including more than 5 components, there is only a modest increase of the explained variance. Therefore, we chose to proceed taking 5 components into account

#### Assessment of the number of clusters in the K-mean clustering (for different kernels of PHATE)

We used three methods to help us define the number of clusters to classify the output of the PHATE algorithm: the elbow, the silhouette, and the gap statistics (see <u>Material and Methods</u>). For each, we ran the PHATE algorithm, with a different number of clusters, followed by the K-means algorithm. Then, we plotted the elbow, silhouette, and gap statistic (supplementary figure <u>SP1</u>). We aimed at observing whether, as the number of kernels and PCA components in PHATE increased, the number of clusters would also increase. The elbow and the silhouette graphical outputs were difficult to interpret and did not yield a clear cutoff for choosing the number of centroids. However, the gap statistic indicated the optimal trade-off at *k*=7 clusters, which was selected for further analyses.

#### Statistical validation

To confirm the result of our analyses, the robustness of the results was assessed through various tests, as shown in figure <u>5</u>. These included 1) examining the informativeness of the clusters as compared to random associations (Fig. <u>5.A</u>), 2) testing the significance of transitions between clusters against random dynamics (Fig. <u>5.D</u>), and evaluating the stability of the clusters across different subjects. Similarly, as already described, the robustness of the dimensionality reduction was verified by comparing the probability of a region being recruited (or not) in a given cluster, to what is observed given random clusters (Fig. <u>5.I</u>) (3).

In figure <u>5.C</u>, it is shown that the entropy of the subject data is significantly lower than the entropy of random clustering. This indicates that the clustering generated by our unsupervised method is more structured than the random clustering. As shown in figure <u>5.B</u> and <u>5.C</u> the null model demonstrates that the initial clusters are unlikely to be due to chance and, instead, reflect the recurrent coordinated activations (de)activations of specific groups of brain regions.

Then, figure <u>5.E</u> shows that the transition probability matrix (containing the probability of sequentially evolving from a state to another state) has higher values in the diagonal, as compared to the off-diagonal transitions However, this probabilities in the diagonal remain lower than what would be expected by chance alone (figure <u>5.H</u> and <u>SP15</u>). Furthermore, there are also preferential transitions between different states (i.e., some sequence of transitions occur more/less often than what is expected by chance). Lastly, according to the entropy plot in figure <u>5.K</u>, we observe that the entropy of our cluster is significantly higher than the entropy distribution of the data in which the regions’ status were shuffled. In other words, the pattern of regional activation/deactivation within the clusters are unlikely to be random. Additionally, it shows that the dimensionality reduction preserves the structure of the manifold throughout the dimensionality reduction.

## Discussion and Conclusion

A converging body of evidence shows that brain activities evolve according to complex, nonlinear dynamics^19^. Measures such as the fractional occupancy, dwell-times and transition probabilities have been used to describe the dynamical properties of the data ^20–22^. These recurrent activation modes are modulated by a number of physiological phenomena^23^ and have been linked to health/disease ^24–27^, task performance^8^, or even consciousness^28^.

In this work, we utilise eyes-closed source-reconstructed MEG scans from 18 healthy young adults during resting-state. Our work builds on recent evidence suggesting that nonlinear transient events, i.e., neuronal avalanches, represent moments where regions interact creating transient configurations, “states”. From this starting point, we could reduce the data complexity through signal binarization, which allowed us to identify transient coordinated events. Then, we applied the PHATE algorithm^12^ to reduce the dynamics to lower dimensionality, and K-means clustering to reduce the number of states by grouping the avalanche patterns.

Neuronal avalanches can be understood as quasi-stable configurations –or attractors– of the dynamics^29^. As the brain’s activity evolves, it alternates between periods of uncoordinated activity, where regions operate independently, and moments of transient coordination, corresponding to neuronal avalanches. Over time, the brain visits multiple attractors (avalanche patterns), resulting in metastable dynamics. Importantly, not all states are visited: at rest, certain regions are more consistently recruited in avalanches (“coordinated”) than others. While brain dynamics is fine-tuned and, as mentioned earlier, its disruptions are related to disease, the specific trajectories (i.e., the sequence in which these states are visited) in the healthy brain remain largely unknown. We hypothesised that the transitions between states follow a regulated (i.e., non-random) sequence of avalanche patterns, with the successive topographies of these recruited patterns exhibiting a distinct structure. To test this hypothesis, we compared our proposed methodology against various surrogate models to demonstrate that the observed patterns are unlikely to have arisen by chance.

Despite neuronal avalanches occurring in less than 5% of the time points (e.g., there are only 3 seconds spent in avalanche activity out of 1 minute recording), these events carry unique information about the brain dynamics. In fact, previous research focusing on the neuronal avalanche statistics, topography, and propagation pathways, showed the potential of these transient events as a marker of healthy brain dynamics^28,30,31^. Thus, focusing on the sequence of avalanche patterns before feeding the data into dimensionality reduction or clustering algorithms might be a neurophysiological-principled way to reduce the computational demand and the noise while focusing on the “most relevant” part of the data. Indeed, in this work, we showed reliable state identification by operating on the neuronal avalanche patterns, hence discarding the rest of the data. In order to achieve this, we employed dimensionality reduction techniques on the sequence of avalanche patterns.

The primary goal of dimensionality reduction is to sift through redundant information in complex, multi-modal datasets without relying on predefined assumptions. This approach maps the effective information contained in the original features to a low dimensional feature space. In this context, the evolution of the activity over time is often described as the system evolving over a low-dimensional surface (i.e., a manifold), rather than exploring the full high-dimensional phase space^32,33^.

When applied to the description of the evolution of neural dynamics, dimensionality reduction techniques like Principal Component Analysis (PCA), Independent Component Analysis (ICA), t-Distributed Stochastic Neighbour Embedding (t-SNE), and Uniform Manifold Approximation and Projection (UMAP) are not explicitly designed to retain both local and global nonlinear relationships in the low-dimensional representation. For instance, t-SNE excels in visualising high-dimensional datasets by minimising the Kullback-Leibler divergence between joint probabilities in the embedding and the original data, but its outcomes can vary due to the usage of gradient descent with random initialization for optimising a non-convex problem^11^.

Hence, these techniques might not be best-suited for data with multiple, nested spatio-temporal scales, such as MEG data. In contrast, as mentioned earlier, PHATE preserves both local and global structures across scales, constructing a diffusion-based informational geometry that effectively captures manifold-intrinsic distances, offering a detailed and scalable low-dimensional embedding, which might be best suited for MEG data. Furthermore, PHATE shows good denoising properties for clearer visualisation, and provides robust, scalable visualisations that capture a broader range of patterns^12^.

Applying PHATE and K-means clustering to MEG avalanche patterns from healthy individuals led to the identification of distinct clusters corresponding to specific brain networks. In particular, based on the cluster identified by the K-means algorithm we computed the transition probability matrix for each subject, capturing the transitions between clusters and also a global transition matrix that aggregates the avalanche patterns across all subjects. After testing the significance of transitions between the clusters, involving shuffled sequences of each subject’s data to serve as a baseline for comparison, we found that the observed transitions are most likely indicative of the population under study and are not random and the transition to the same cluster is the most probable transition meaning that the system possesses a form of memory about the previously recruited pattern.

The comparison with more standard PCA reduction (figure <u>3.A</u>, <u>3.B</u> and <u>3.C</u>), demonstrates the effectiveness of PHATE in capturing the flow of neural activity, enhancing both the cluster identification and the visualisation of brain states. This approach was further validated through sensibility analysis and robustness testing, confirming the reliability of the findings (sections <u>Sensibility analysis</u> and <u>Statistical validation</u>).

Dimensionality reduction techniques applied to resting state data suffer, in general, from the lack of validation about the “goodness” of the clusters. To this end, we devised a large array of null-models to extensively test both the spatial structure of the clusters, which is unlikely to emerge given a random clustering of the regions, and their evolution, which is not justified by a random sequence of the spatial clusters.

To our knowledge, this is the first study applying the PHATE algorithm to MEG data. In particular, the algorithm had been already applied to raw EEG^34^ and fMRI data^35–38^, where it is shown that PHATE could visualise brain states and help identify altered brain dynamics in early stages of schizophrenia. While these studies have provided convincing evidence of the efficacy of PHATE (e.g., ^38^ with the T-PHATE algorithm or ^36^ combining data from several sources, including fMRI), none of the previous works focused on neuronal avalanches (in MEG data).

It would nonetheless be interesting to further investigate limitations of the PHATE algorithm regarding auto-correlating signals such as fMRI^35^. Indeed, in such data, we may want to explore an approach combining PHATE with a model of signal autocorrelation. In addition, a study presents another limitation of PHATE algorithm^39^, as discrete structures could be more accurately defined, although it also underlines PHATE’s capability in preserving graph distances and local and global ones, assuming multiple tests of parameters are realised.

In the end our study highlights the potential for such dimensionality reduction for MEG data as well as the potential outputs that can be obtained from this. In that regard, further studies on MEG data may benefit from using the PHATE algorithm.

Our findings also reveal well-defined activation patterns, temporality and spatially distinct, expressed in terms of transient coordinated events, in the brain activity among healthy subjects. The clusters appear to be biologically meaningful, in that they either correspond to well-defined anatomical structures (e.g., the left and right frontal lobes in figure <u>4</u>), or they are reminiscent of known functional clusters, such as the default mode network. Crucially, if one performs dimensionality reduction directly from the raw data (i.e., not from the avalanche patterns), the algorithm fails to retrieve meaningful, non-random patterns. In other words, the dynamics of spontaneous activities, based on avalanche patterns, is constrained to prefer specific sequences and to go across other sequences less often than expected (given the relative dwell-time across state, which is kept constant in the surrogates). Such constraints are preserved in the low dimensional representations. In conclusion, our work demonstrates that brain dynamics can be understood as a fine-tuned sequence of intermittent nonlinear transient coordinated events (neuronal avalanches) that modify their spatio-temporal structure over time. The implications are manifolds, as nonlinear transient events are typically overlooked in favour of a more oscillatory perspective. This approach allows a theory-driven, mathematically-principled way to vastly reduce the complexity of the data. After this step, the deployment of advanced dimensionality-reduction techniques successfully led to a parsimonious data-driven description of the large-scale dynamics.

## Acknowledgements

The work was supported by the European Union’s Horizon 2020 research and innovation program under grant agreement No. 101147319 (EBRAINS 2.0 Project) and No. 101137289 (Virtual Brain Twin Project) and by the EBRAINS Italy nodo Italiano grant, CUP B51E22000150006.

## Author Contributions

A.E.C., L.K., G.R., P.S. Conceptualization and Methodology;

A.E.C., L.K., M.A., G.R., P.S. Simulations and Data analysis;

A.E.C., L.K., M.A., G.R., P.S. Formal analysis;

J.R., R.C., A.E., V.J., P.S. Resources;

A.E.C., L.K., M.A., G.R., P.S. Writing - Original Draft;

A.E.C., L.K., M.A., E.T.L., A.P., R.M., A.R., V.J., G.R., P.S. Writing - Review - Editing;

A.E.C., L.K., M.A., E.T.L., A.P., R.M., A.R., V.J., G.R., P.S.Visualization;

V.J., G.R., P.S. Supervision and Project administration;

V.J., P.S. Funding acquisition;

All authors read and approved the final manuscript.

## Data availability

The data cannot be made public due to its personal nature, and as this was foreseen in the informed consent.

## Code availability

The code is openly and publicly accessible at: https://github.com/lionelkusch/patien-analysis.

## Abbreviations

AAL: Automated Anatomical Labelling
ECG: Electrocardiogram
EEG: Electroencephalography
EOG: Electrooculogram
fMRI: Functional Magnetic Resonance Imaging
ICA: Independent Component Analysis
MEG: Magnetoencephalography
MRI: Magnetic Resonance Imaging
PCA: Principal Component Analysis
PHATE: Potential of Heat-diffusion for Affinity-based Transition Embedding
SD: Standard deviation
TE: Echo time
TI: Inversion time
TR: Repetition time
t-SNE: t-Distributed Stochastic Neighbour Embedding
UMAP: Uniform Manifold Approximation and Projection

## Ethical statement

The study was approved by the Ethics Committee ASL-NA1 centro (Prot.n.93C.E./Reg. n.14-17OSS), and all participants provided written informed consent.

## Supplementary figures

**Fig SP1.**
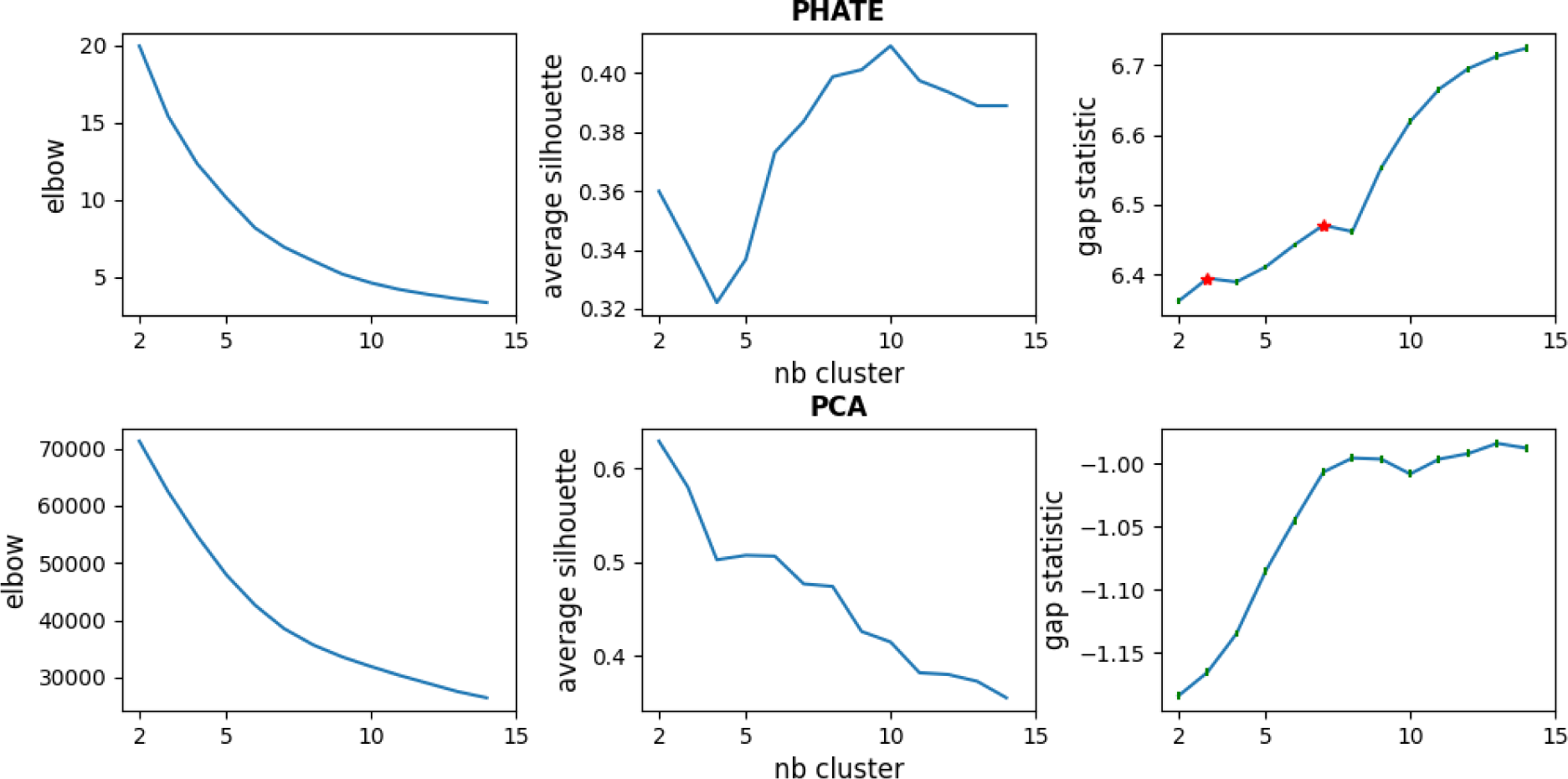
Compare number of clustering of Phate and PCA. The top and bottom panels respectively evaluate the number of clusters of low dimension of PHATE and PCA.

**Fig SP2.**
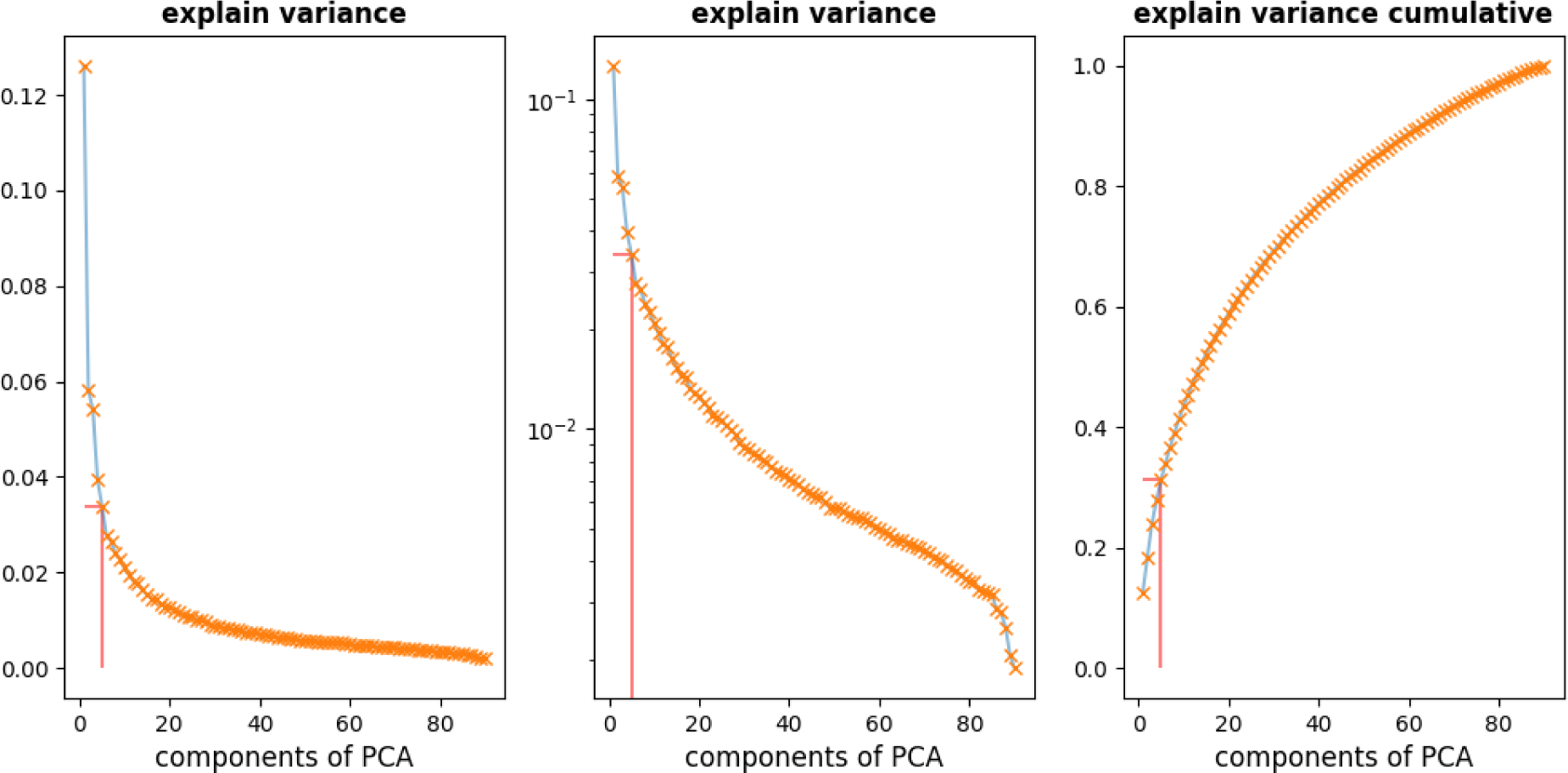
Compare number explained variance by number of components and cumulative explained variance for PCA. This figure displays PCA explained variance by component (left and centre) and PCA cumulative explain variance (right).

**Fig SP3.**
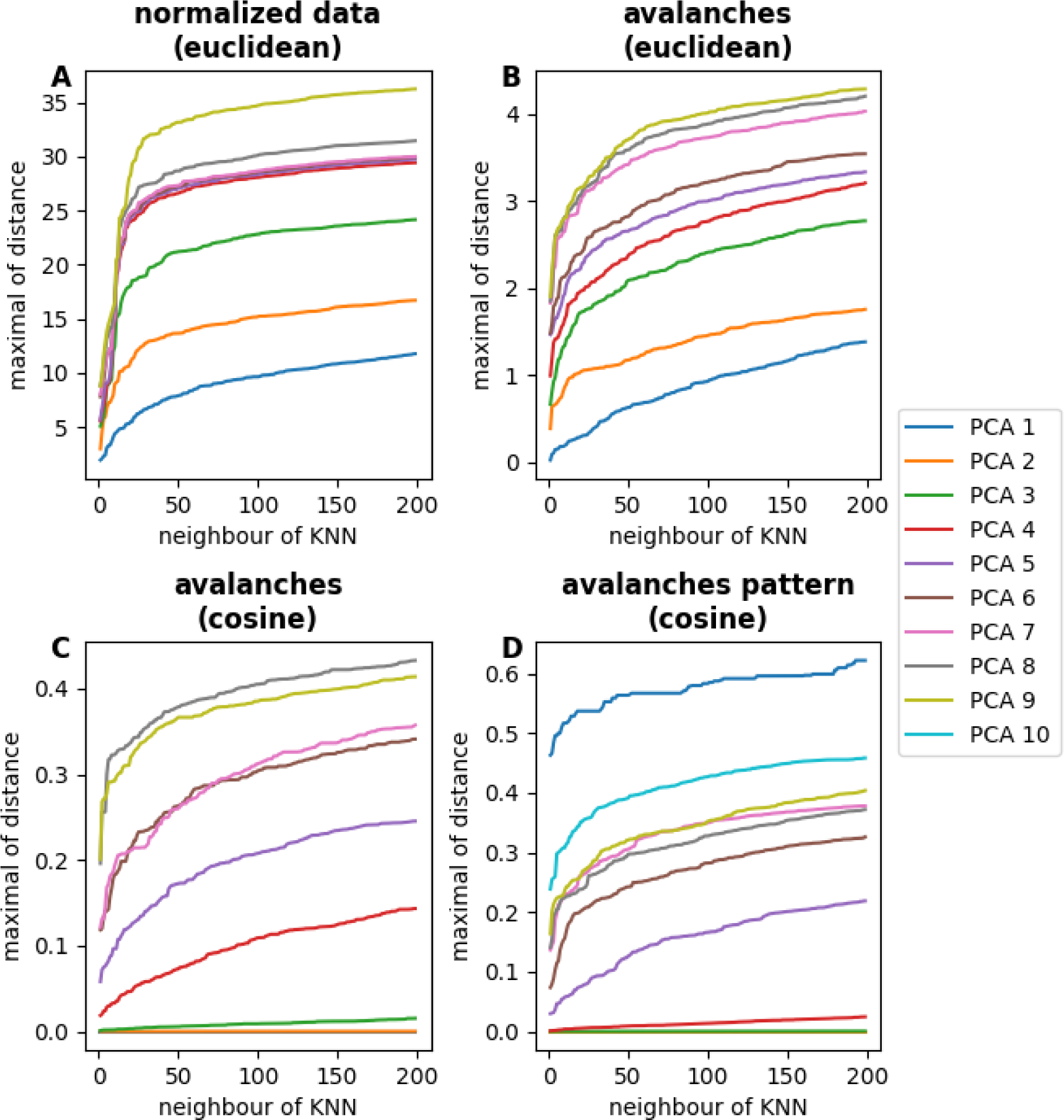
Distance of KNN in PCA for different numbers of components for the different steps of the pipeline. This figure evaluates the maximal distance between k-neighbours in KNN based on euclidean distance for normalised data (**A**) and avalanches (**B**) and on cosine distance for avalanches (**C**) and avalanches pattern (**D**). Each line corresponds to a number of PCA components (2 to 15).

**Fig SP4.**
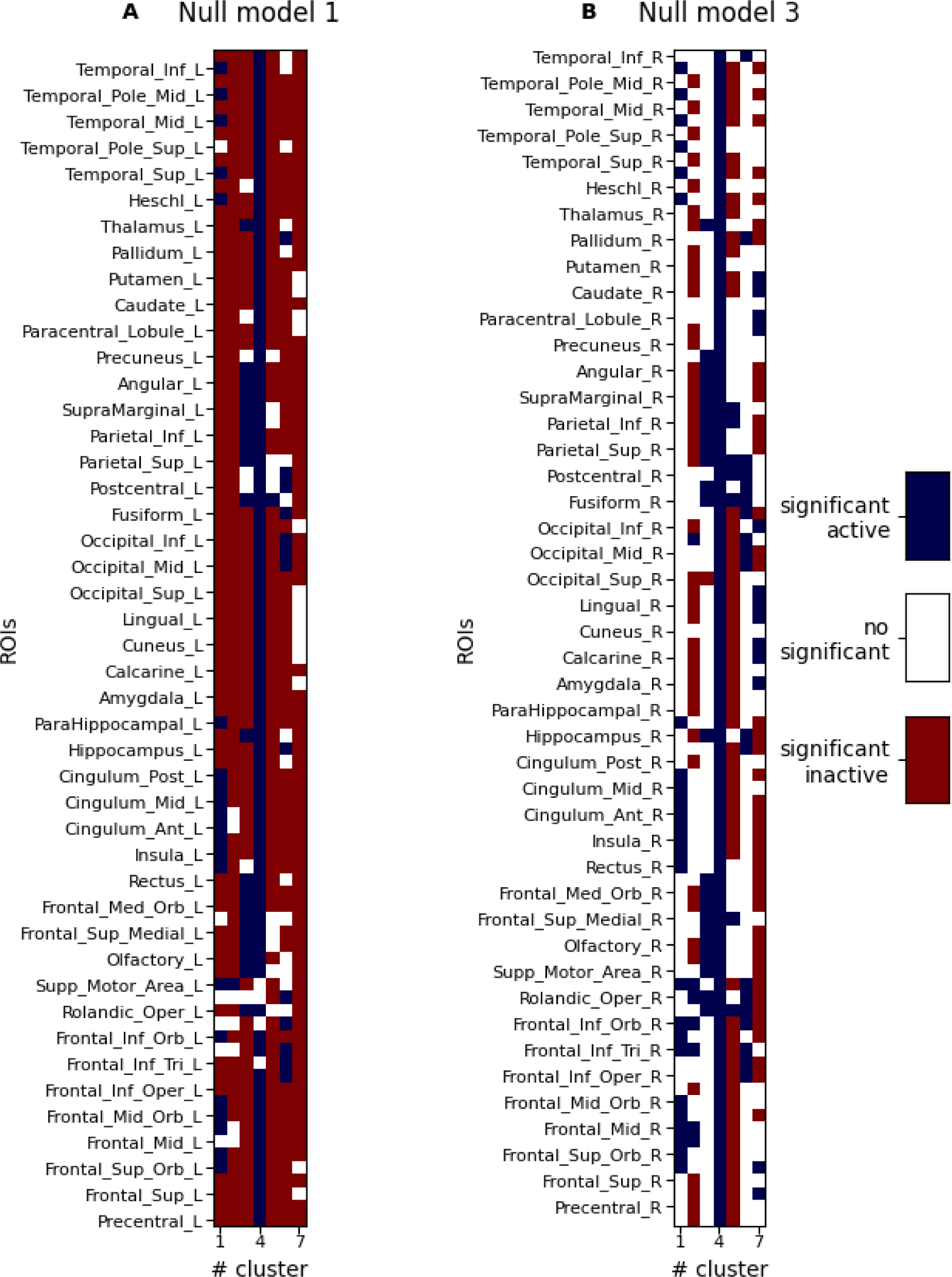
Significant of brain region activity in each cluster. Each matrix from **A** and **B** displays the significant vectors of all clusters. In other words, they represent the sum of avalanches belonging to the cluster, resulting from the association of each avalanche to a specific cluster. More specifically, those panels show the significance of the probabilities of an avalanche activation (red), deactivation (blue) or at basal state (white) to be part of a cluster, for a precision at 0.05 (**A**) and 0.2 (**B**).

**Fig SP5.**
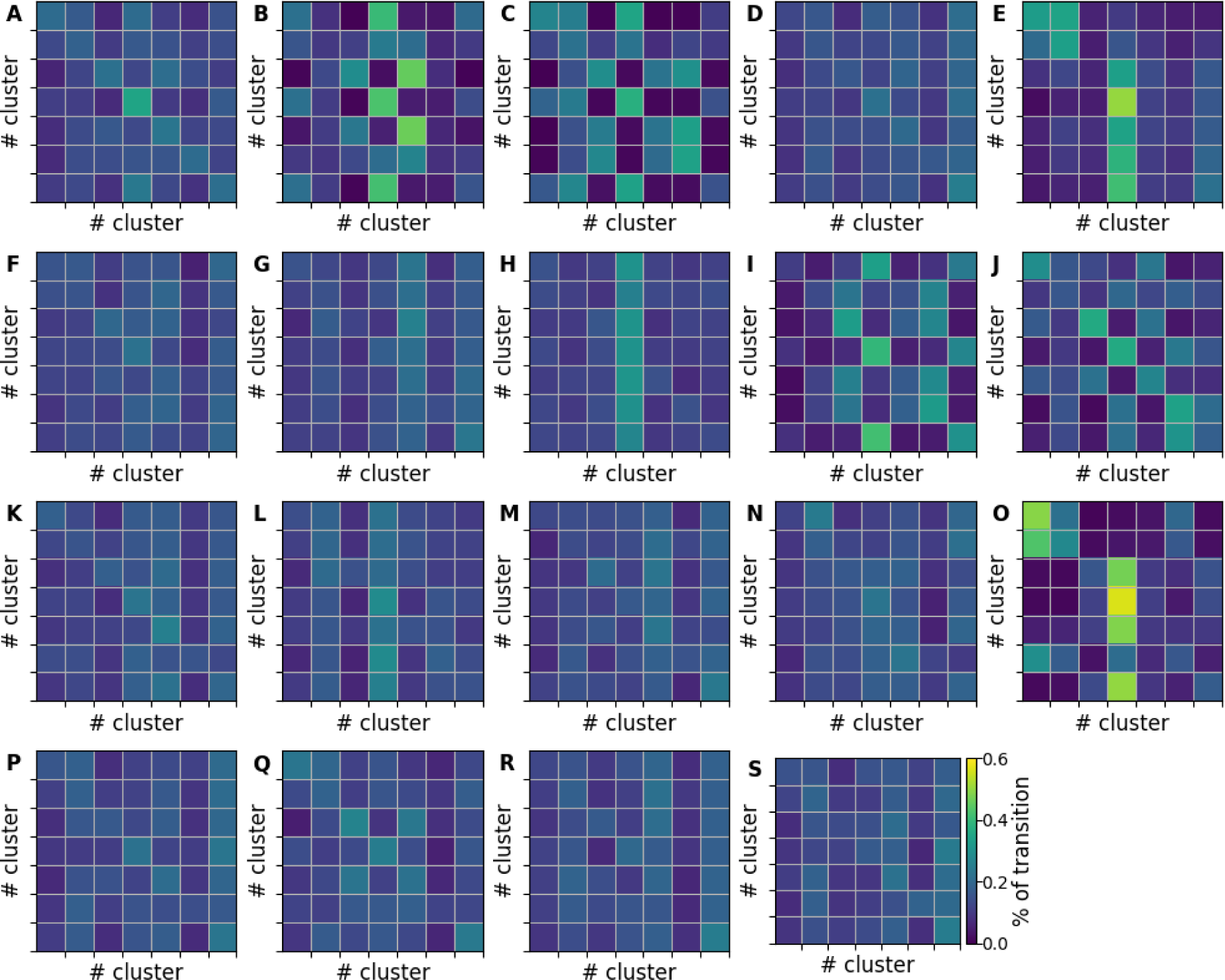
Transition probability matrix for all subjects and for each one. The top left panel (**A)** displays the transition probabilities matrix for all subjects concatenated. The others (**B-S**) represent the transition probabilities matrices for each subject.

**Fig SP6.**
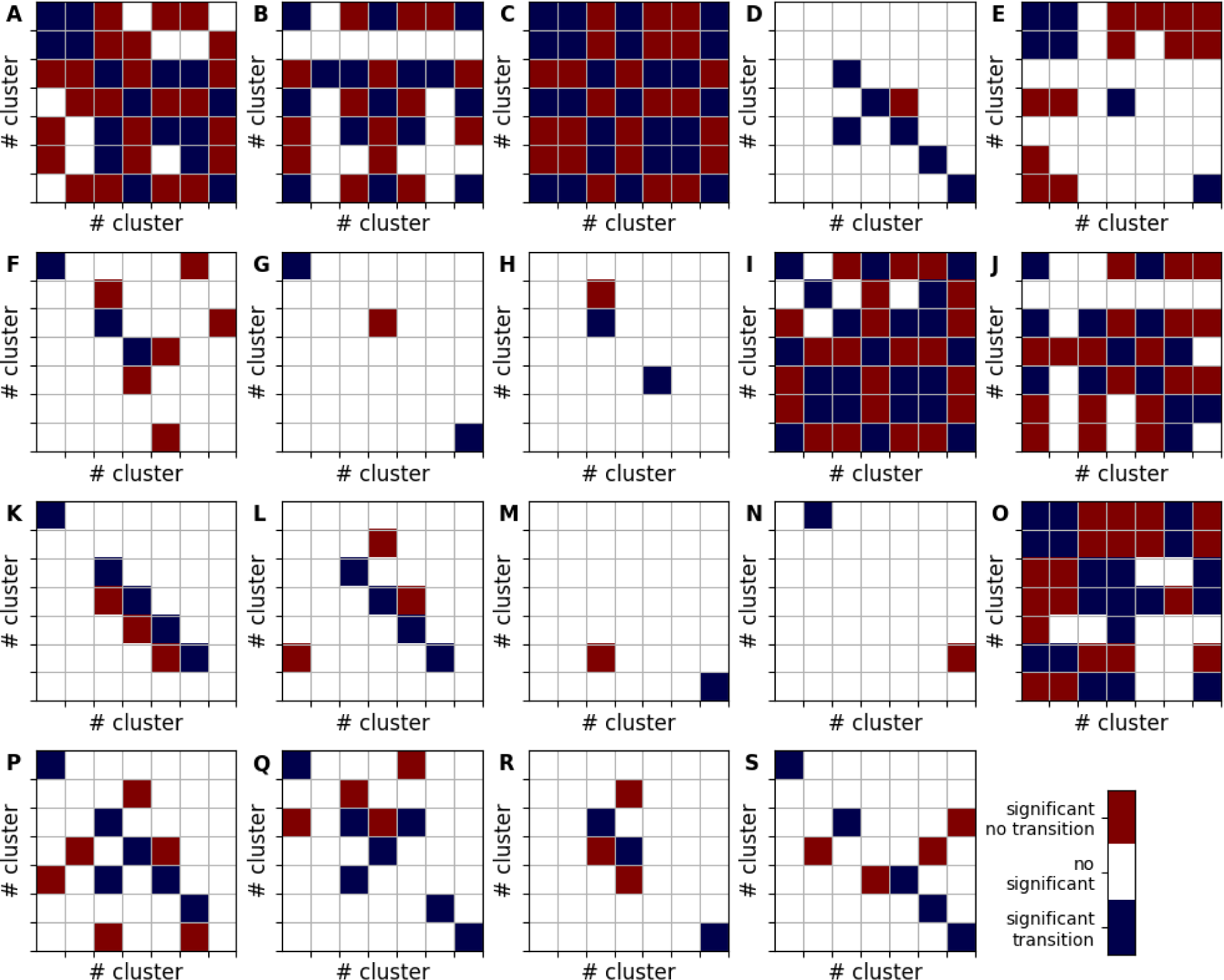
Significant transitions matrices for all subjects and for each one. The top left panel (**A)** represents the significant transitions for all subjects concatenated. The others (**B-S**) represent the significant transitions for each subject. The precision is set at 0.05.

**Fig SP7.**
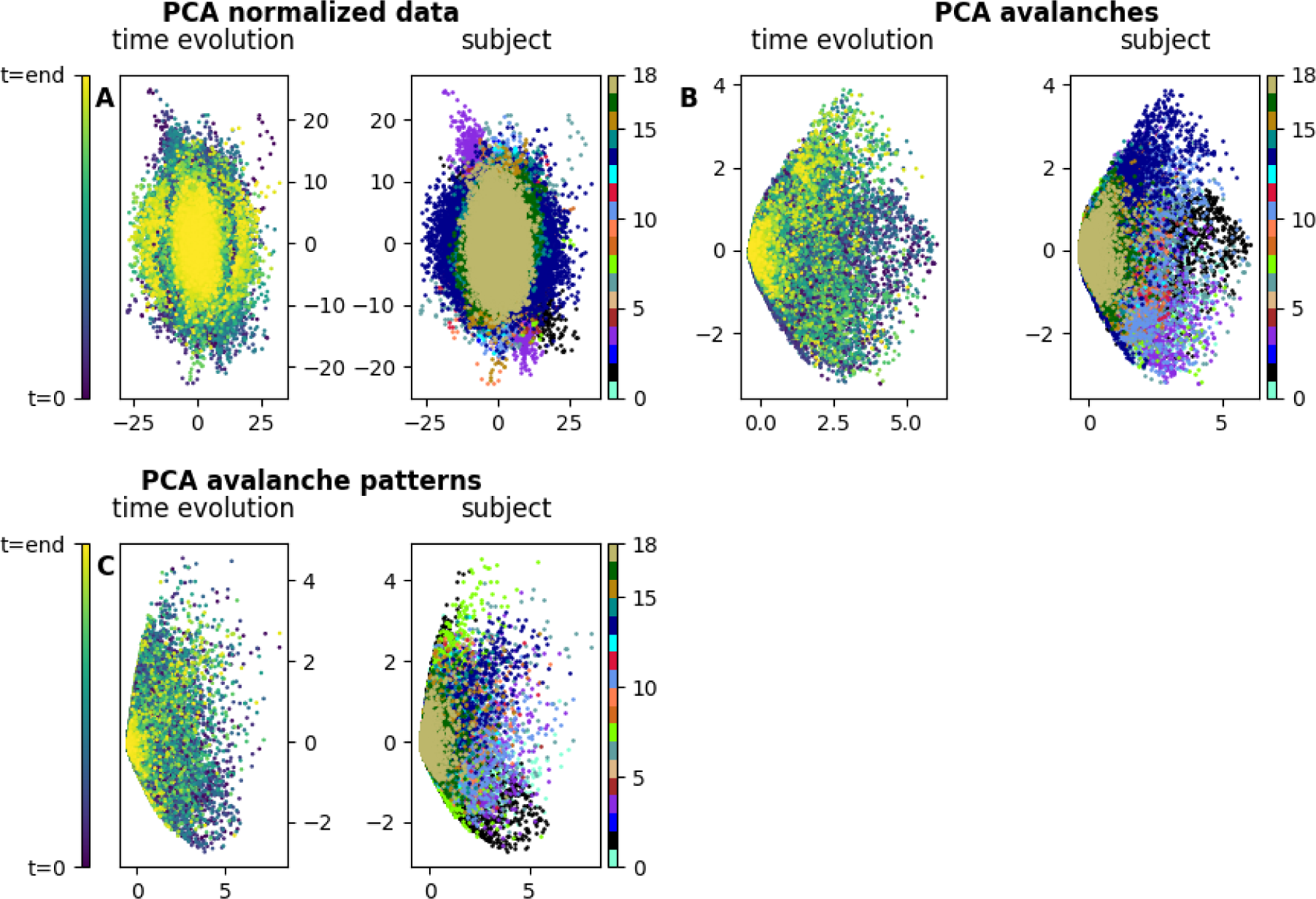
Comparison of PCA results from the different steps of the pipeline according to time or per subject. The results of PCA for the different steps of the pipeline (**A**: reconstructed z-scored Meg source signal, **B**: binarized activity and **C**: avalanches pattern) compares the time evolution of brain state (left, time is the colour code) and subject data (right, subject is the colour code) for the subjects in the two first components of the PCA space.

**Fig SP8.**
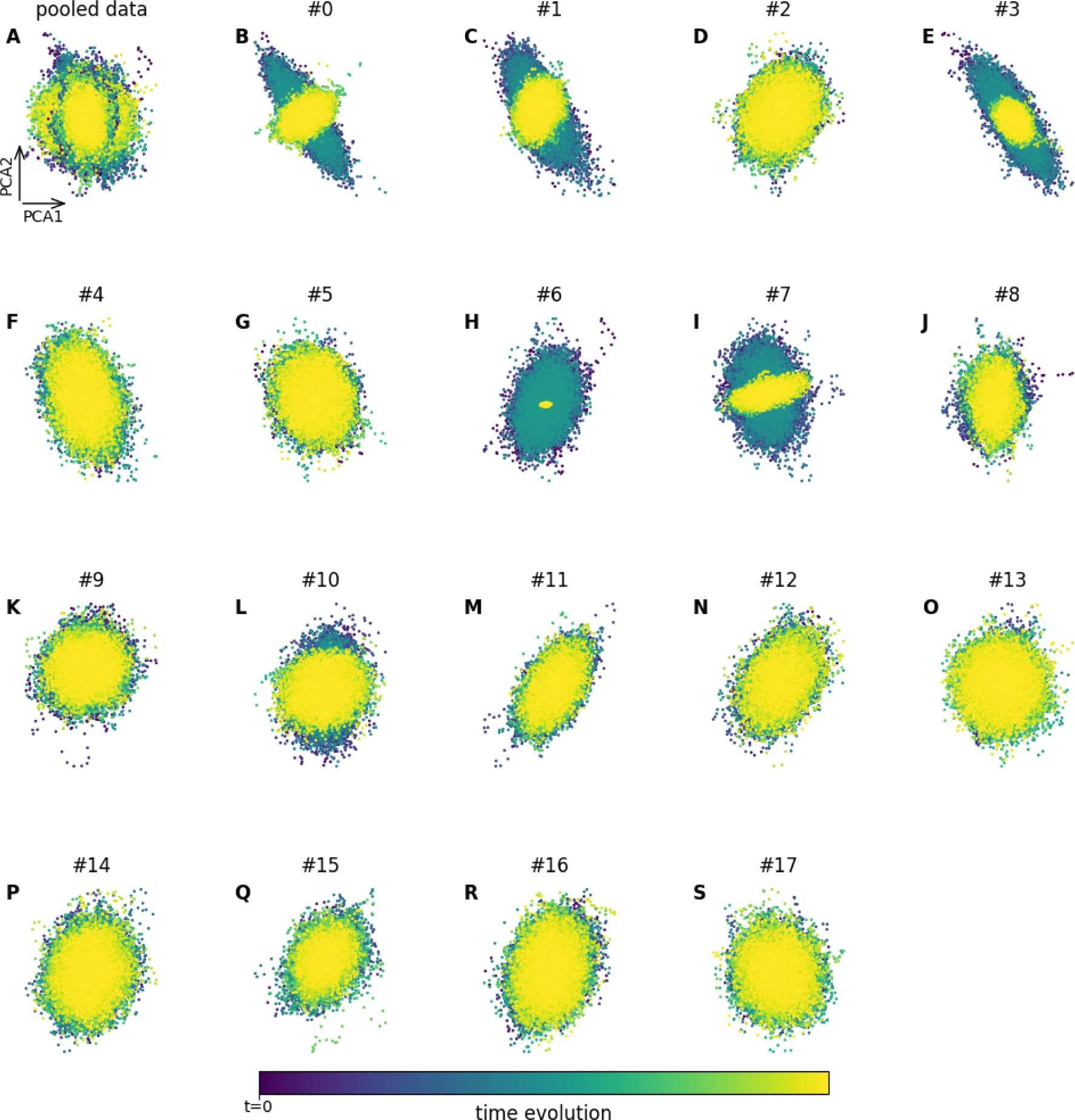
PCA on reconstructed z-scored MEG source signals. The graphic shows the time evolution of brain state for the subjects in the two first components of the PCA space. The top left panel (**A**) corresponds to a superposition of all subjects. The other graphs (panels **B-S**) are for each subject. Hence, in these plots, each dot, each dot represents the evolution of the whole system brain state at each instant (colour code).

**Fig SP9.**
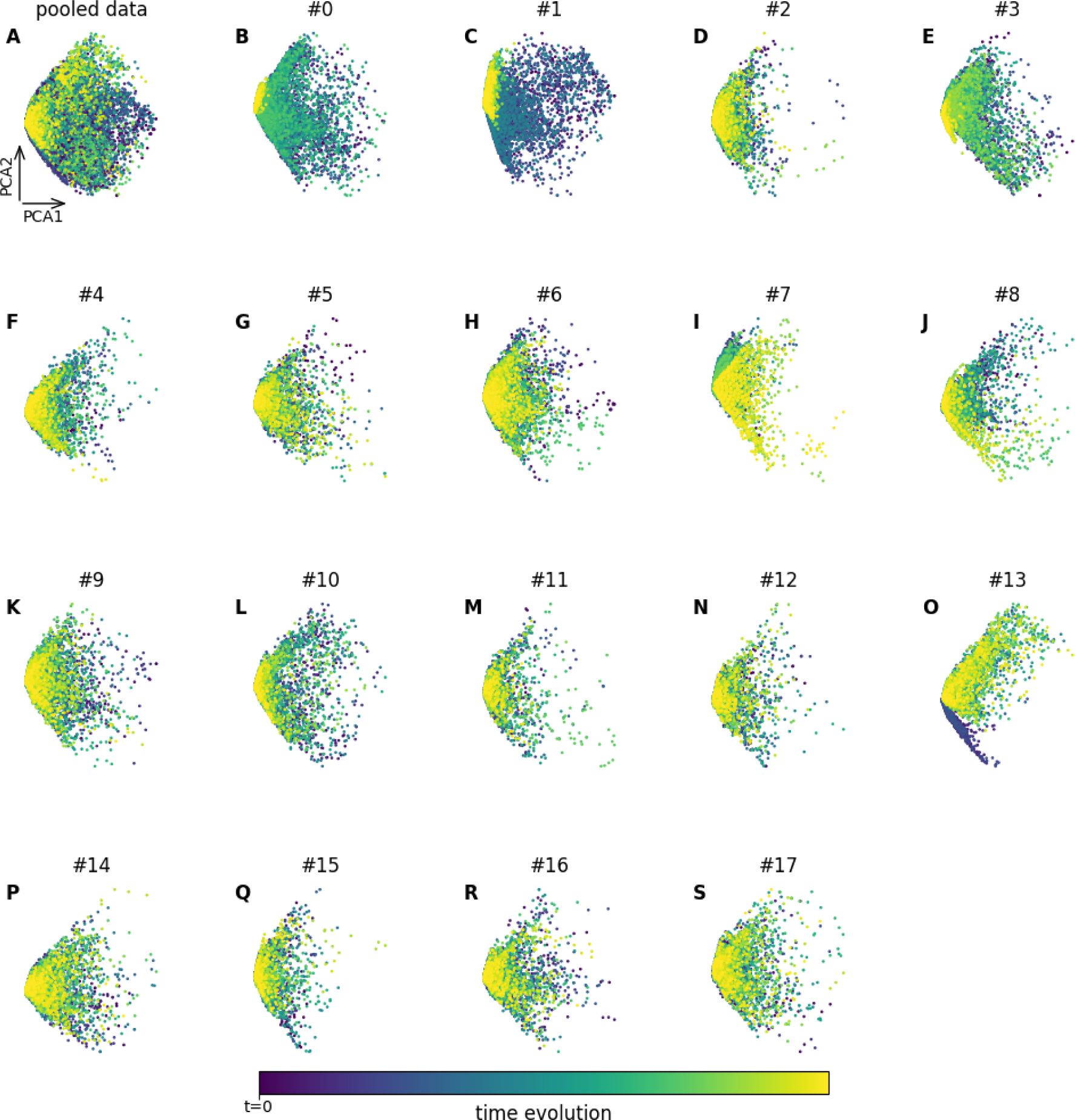
PCA on avalanches. The graphic shows the time evolution of brain state for the subjects in the two first components of the PCA space. The top left panel (**A**) corresponds to a superposition of all subjects. The other graphs (panels **B-S**) are for each subject. In these plots, each dot, each dot represents the evolution of the whole system brain state at each instant (colour code).

**Fig SP10.**
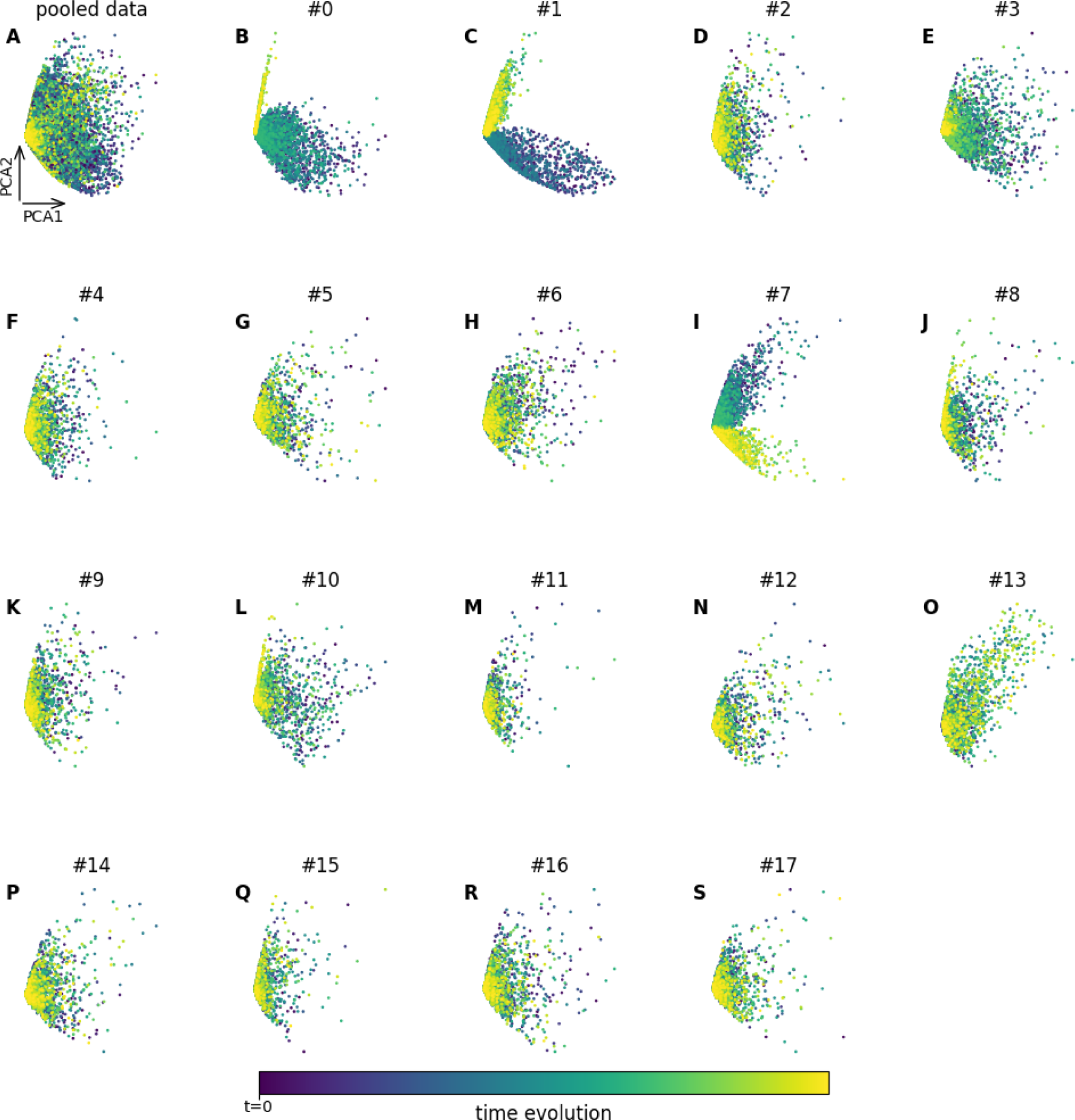
PCA on avalanches patterns. The graphic shows the time evolution of brain state for the subjects in the two first components of the PCA space. The top left panel (**A**) corresponds to a superposition of all subjects. The other graphs (panels **B-S**) are for each subject. In these plots, each dot, each dot represents the evolution of the whole system brain state at each instant (colour code).

**Fig SP11.**
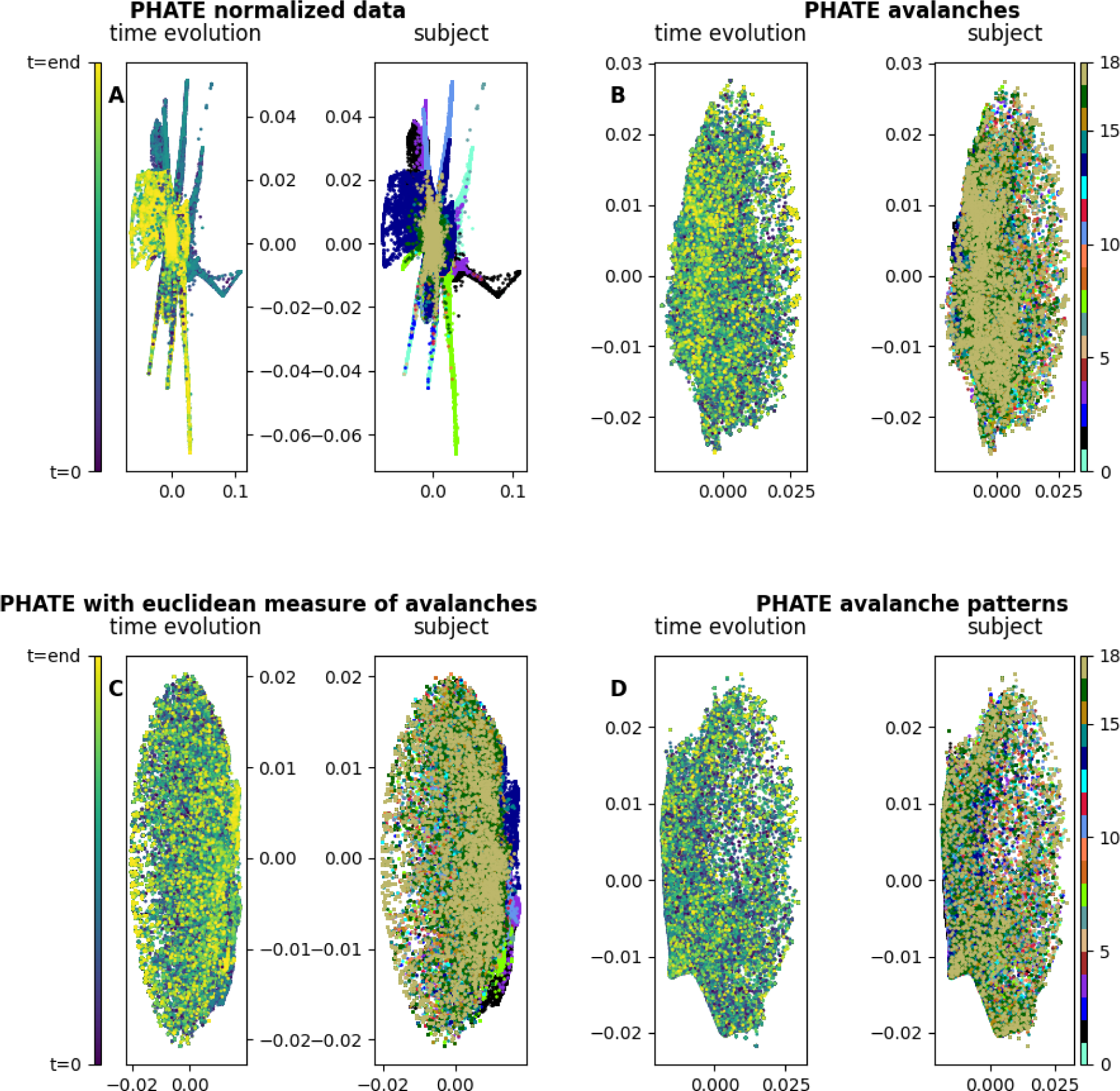
Comparison of PHATE results from the different steps of the pipeline. The result of PHATE for the different steps of the pipeline (**A**: z-scored signal, **B**,**C**: binarized activity and **C**: avalanches pattern) shows the time evolution of brain state (left, time is the colour code) and subject data (right, subject is the colour code) for the subjects in the two first components of the PHATE space.

**Fig SP12.**
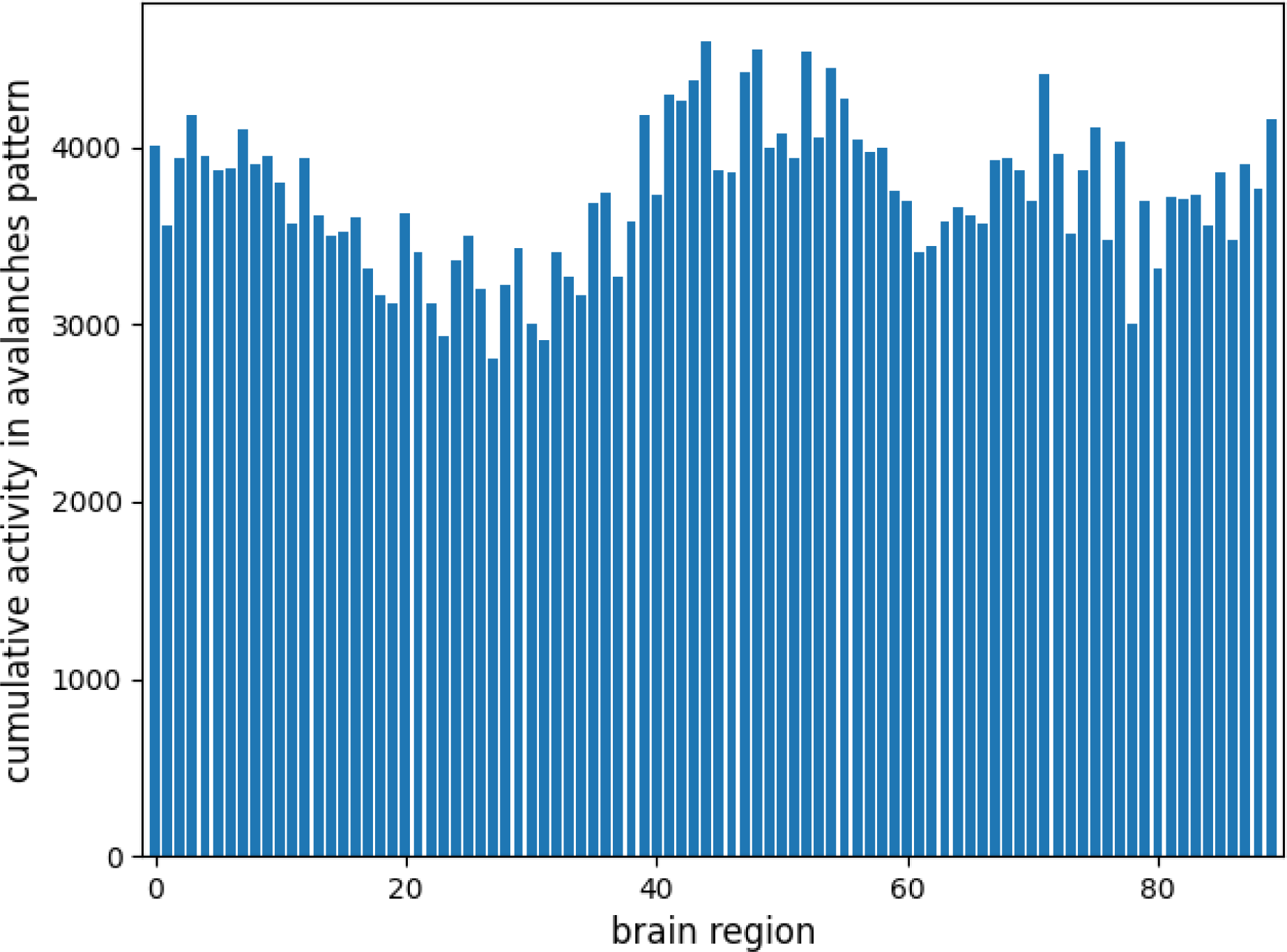
The cumulative of activation in all avalanche patterns. The figure represents the cumulative activation of each region in avalanches pattern.

**Fig SP13.**
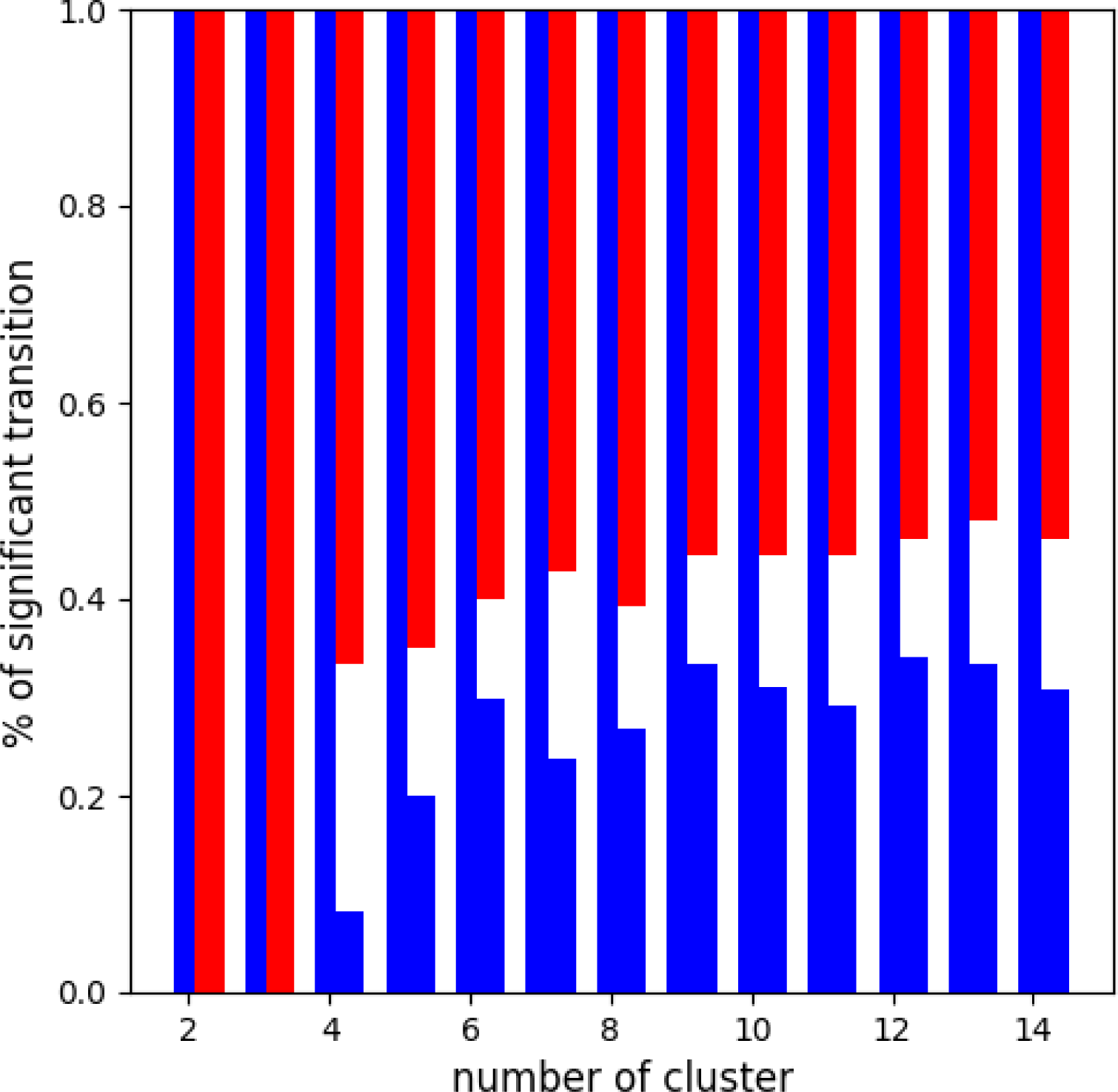
Stability of the clusters. This panel displays two columns by different numbers of clusters. The left column corresponds to the percentage of the diagonal matrix of the transition between clusters that is higher or lower than chance and the left column is for the rest of the transition. The blue indicates the percentage of transitions which are lower than chance and the red, the percentage of transitions which are higher than chance. The significance threshold is set at 0.05.

**Fig SP14.**
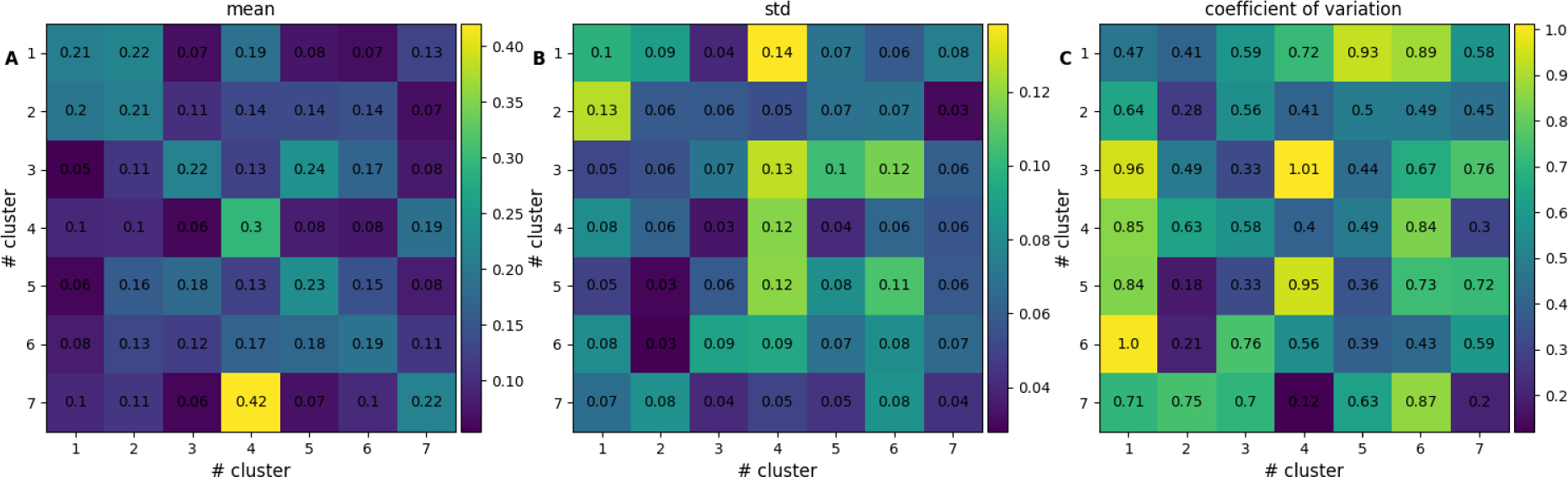
Variability of the significant transition between clusters. The graphics represent the mean (**A**), standard deviation (**B**) and the coefficient of variation (**C**) of the transition of all subjects.

**Fig SP15.**
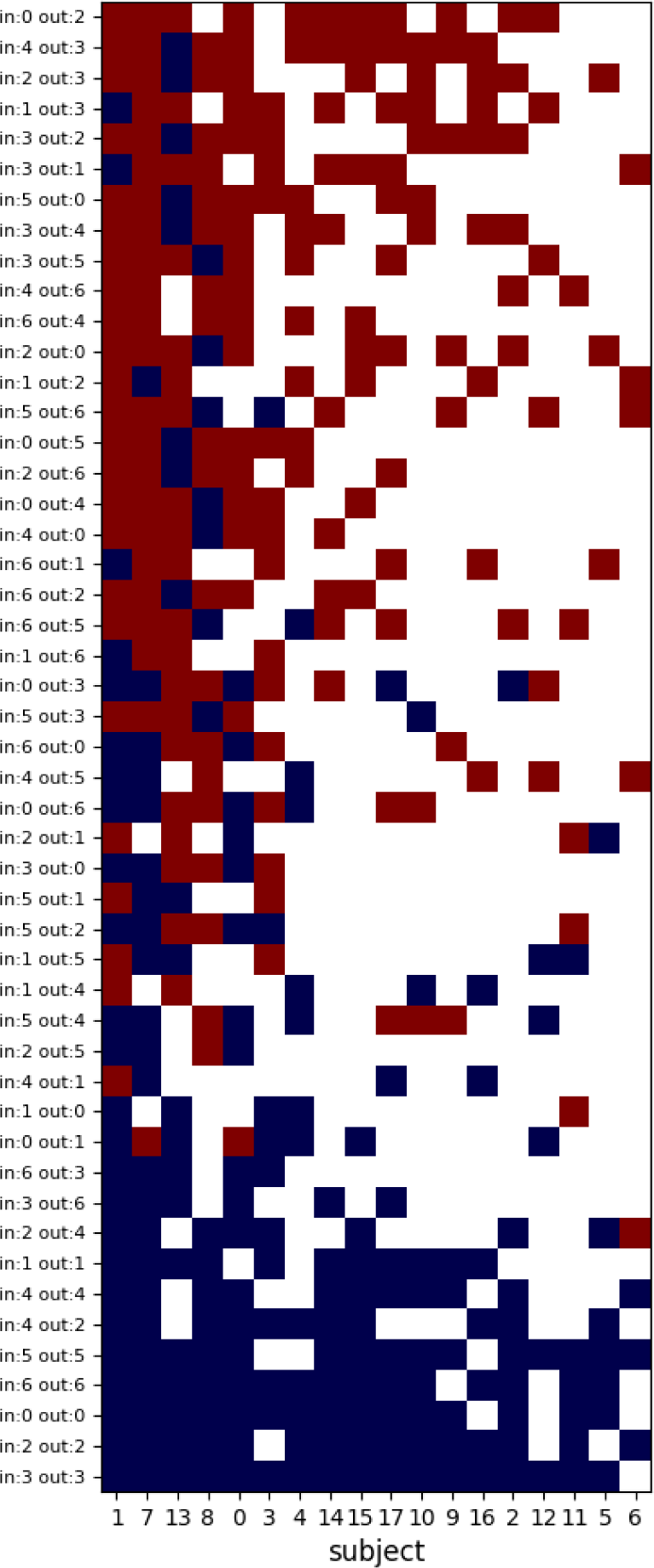
Comparing the significant transition between cluster in each subject. This panel orders the transition between clusters from the cluster with higher transition of probability than the chance to lower transition probability. The subjects are ordered in function of their number of significant transitions.

